# A suite of selective pressures supports the maintenance of alleles of a *Drosophila* immune peptide

**DOI:** 10.1101/2023.08.18.553899

**Authors:** Sarah R. Mullinax, Andrea M. Darby, Anjali Gupta, Patrick Chan, Brittny R. Smith, Robert L. Unckless

## Abstract

The innate immune system provides hosts with a crucial first line of defense against pathogens. While immune genes are often among the fastest evolving genes in the genome, in *Drosophila*, antimicrobial peptides (AMPs) are notable exceptions. Instead, AMPs may be under balancing selection, such that over evolutionary timescales multiple alleles are maintained in populations. In this study, we focus on the *Drosophila* antimicrobial peptide Diptericin A, which has a segregating amino acid polymorphism associated with differential survival after infection with the Gram-negative bacteria *Providencia rettgeri*. Diptericin A also helps control opportunistic gut infections by common Drosophila gut microbes, especially those of *Lactobacillus plantarum*. In addition to genotypic effects on gut immunity, we also see strong sex-specific effects that are most prominent in flies without functional *diptericin A*. To further characterize differences in microbiomes between different *diptericin* genotypes, we used 16S metagenomics to look at the microbiome composition. We used both lab reared and wild caught flies for our sequencing and looked at overall composition as well as the differential abundance of individual bacterial families. Overall, we find flies that are homozygous for one allele of *diptericin A* are better equipped to survive a systemic infection from *P. rettgeri*, but in general have a shorter lifespans after being fed common gut commensals. Our results suggest a possible mechanism for the maintenance of genetic variation of *diptericin A* through the complex interactions of sex, systemic immunity, and the maintenance of the gut microbiome.

## Introduction

An effective immune response is essential for organisms to protect themselves from pathogens. However, an excessive immune response causes damage to self either directly (autoimmunity, cytokine storms) or through dysbiosis *via* disruption of the composition of beneficial microflora [1–4]. The dynamic suite of microbes that hosts face necessitates initiating a robust systemic immune response, avoiding self-harm, and maintaining a beneficial microbiome. The challenge of maintaining a balanced immune response is exacerbated because the threat of microorganisms is contextual; thus, the immune system must distinguish harmful versus beneficial microbes in each context [5,6]. For example, an ingested microbe may be harmless or even beneficial as a food source in the digestive tract, but, when in circulation, that same microbe may prove harmful to the organism [7]. One consequence of balancing a robust innate immune system is that it may lead to the maintenance of genetic polymorphism in genes that encode for proteins intimately involved in the immune response [8,9]. Such patterns of maintained polymorphisms contrast the standard coevolutionary arms race model where hosts and pathogen continually adapt to each other leading to rapid evolutionary change in immune genes [10–12]. Though the coevolutionary arms race model appears apt in many cases, the case for the adaptive maintenance of alleles, balancing selection in the broad sense, on genes involved in the immune defense stems from a more nuanced view of the delicate interplay of systemic immunity, life history traits and the beneficial microbiome [13,14].

Balancing selection is the general term for the adaptive maintenance of multiple alleles in a population. In contrast to the evolutionary arms race model, natural variation in many immune genes is maintained by balancing selection [15–18]. The inference of balancing selection from molecular population genetic data is difficult, especially when genes are small and in areas of high recombination. Nonetheless, several examples of balancing selection exist, many of which involve small effector proteins of the innate immune system [18–21].

Balancing selection on immune genes is likely to involve allelic benefits that are only conditionally beneficial. One way that an allele could be conditionally beneficial is if there is specificity between pathogen and allele [22] such that *allele A* better protects against *pathogen 1* and *allele B* better protects against *pathogen 2*: *pathogen specificity hypothesis*. Another way that an allele can be conditionally beneficial in the context of immune defense is if resistance alleles are costly in the absence of infection. In this case, *allele A* better protects against both *pathogen 1* and *pathogen 2*, but in the absence of infection *allele A* is costly for its host [23]. This cost could be energetic or through autoimmune-like damage: *autoimmune hypothesis*. In reality, there is likely a continuum between these alternative hypotheses.

Invertebrates lack an adaptive immune system, so the delicate balance between systemic and gut immunity is achieved only through adjustments to the innate immune response [24]. It is therefore important to understand how invertebrates optimize their systemic immune response with as little detriment to their beneficial gut microbiota as possible.

Though first characterized for its role in systemic immunity, the (IMD) pathway is also the main NF-κB immune pathway in the gut and contributes to the maintenance of the microbiome and protection from gut infections [25–27]. To date, there is little research into how individual AMPs help maintain a healthy microbiome composition in the fly, let alone how allelic variation may affect microbiome composition [28,29]. The standard gut microbiome of lab reared *D. melanogaster* consists of two prominent bacterial genera: *Lactobacillus* and *Acetobacter* [27]. Both genera are easily cultured in the lab, and the *Drosophila* gut can easily be manipulated, making it a good model to study the effects of immune gene variation on microbiome composition and how microbiome composition can, in turn, influence host fitness.

Antimicrobial peptides (AMPs) are a critical part of the innate immune system that act as broad-spectrum antimicrobials, combating bacteria, fungi, and viruses. In the coevolutionary arms race model, AMPs are on the front lines: directly interacting with microbes. AMPs have also been demonstrated to have critical roles in other physiological functions, for example, dysregulation of AMPs has been connected to diseases such as atopic dermatitis and Alzheimer’s disease [2,30,31]. AMPs also contribute to the aging process where the immune system becomes more active as organisms ages to compensate for a decline in its effectiveness. However, this leads to cytotoxicity that in turn shortens lifespan [3].

In *Drosophila,* AMPs play a crucial in both manage systemic infections and in the maintenance of the gut microbiota [32]. AMPs, however, appear to evolve more slowly than most immune gene families [33–36]. One explanation for this perceived lack of adaptive evolution is that *Drosophila* AMPs genetic variation is maintained adaptively through balancing selection [15,37–39], where different selective pressures favor different alleles. Naturally occurring allelic variation in AMP loci are sometimes associated with variability in pathogen resistance [19,40–42]. *Diptericin A* (hereon referred to as *diptericin* or *Dpt)* is one of the canonical effector genes of the *Drosophila* Immune deficiency (IMD) pathway, and is generally associated with defense against Gram-negative bacterial infection [43,44].

In both *D. melanogaster* and *D. simulans*, an amino acid polymorphism at the 69^th^ residue of the mature 83-residue Diptericin peptide segregates in most populations surveyed [19]. In both species, this polymorphism is due to a point mutation that changes the ancestral serine (S) allele to arginine (R), but the two species use different codons for arginine. Unckless *et al.* [19] found a significant difference in survival based on *diptericin* genotype after systemic infection with the Gram-negative bacteria, *Providencia rettgeri,* a natural pathogen of *D. melanogaster.* The study used inbred fly lines and found homozygous serine flies have a better survival rate 5 days post infection (∼60%) than homozygous arginine flies (∼20%) or flies with a premature stop codon in *diptericin* (0%). Later work by Hanson *et al.* took a more general approach to AMP specificity and found that *Diptericin* plays a disproportionate role in response to infection with *P. rettgeri* [45]. What remains unclear, however is why this presumably deleterious arginine allele persist in populations of two different species. We hypothesize that it either protects against a different suite of pathogens or it is beneficial in the absence of infection through some life history tradeoff.

This study aimed to test two hypotheses about the maintenance of genetic variation in *Diptericin.* First, different *Diptericin* alleles might protect against different pathogens – we refer to this as the *pathogen specificity hypothesis*. It would be supported if the arginine allele were to be associated with higher survival than the serine allele in some infections, since the serine allele is already clearly more protective against *P. rettgeri* [19,42]. In contrast, flies with the serine allele might be generally better at surviving infection, but this may have a cost to the organism. We refer to this as the *autoimmune hypothesis,* <u>where a protective immune allele is associated with a tradeoff that is costly in the absence of infection.</u> We test whether there are tradeoffs between systemic immunity and other life history traits – particularly traits related to the gut microbiome utilizing genetically controlled lines for *Diptericin* alleles generated by CRISPR/Cas9-editing. Flies were systemically infected with a panel of six different bacteria and homozygous serine flies had a better 5-day survival with each bacterium. We then used axenic and gnotobiotic flies to look at the lifespan of flies since AMP overexpression and microbial proliferation are commonly observed in aging flies [46–49]. *Lactobacillus plantarum* is harmful to female flies with non-functional diptericin, while homozygous arginine flies poly-associated with *L. plantarum* and *A. tropicalis* had a longer lifespan than homozygous serine flies. In this way we found evidence for a tradeoff between the ability to fight systemic infection and the ability to control opportunistic gut infections.

## Results

### A single amino acid change drastically influences survival after infection

Unckless et al. [19] found that in inbred lines from the *Drosophila Genetic Reference Panel* (DGRP), flies homozygous for serine as position 69 of the mature Diptericin peptide survive systemic infection with *P. rettgeri* much better than those homozygous for the arginine peptide at the same position. This experiment showed only an association between immune defense and the serine/arginine polymorphism since these inbred lines were on several different genetic backgrounds. To control genetic background, we used CRISPR/Cas9 genome editing to create both an arginine allele (single nucleotide change, *dpt^S69R^*) and multiple null alleles (1 or 3 base pair deletions, *Δdpt* flies_)_, as well as control *dpt^S69^* (serine at position 69 of the mature peptide) in *diptericin* (Fig S1A, Table S1). The phenotype for our CRISPR/Cas9 edited flies showed striking similarity to the inbred lines. In systemic infection challenges with *P. rettgeri, dpt^S69^* flies are better protected from infection than *dpt^S69R^* flies (p= 5.42e-08) and *Δdpt* flies (p=6.46e-09, Fig 1A). Remarkably, for inbred lines and CRISPR/Cas9 edited lines, survival for five days post infection with *P. rettgeri* for the serine allele lines is 50-60%, with the arginine allele is 10-20%, and less than 5% for null alleles.

**Figure 1:**
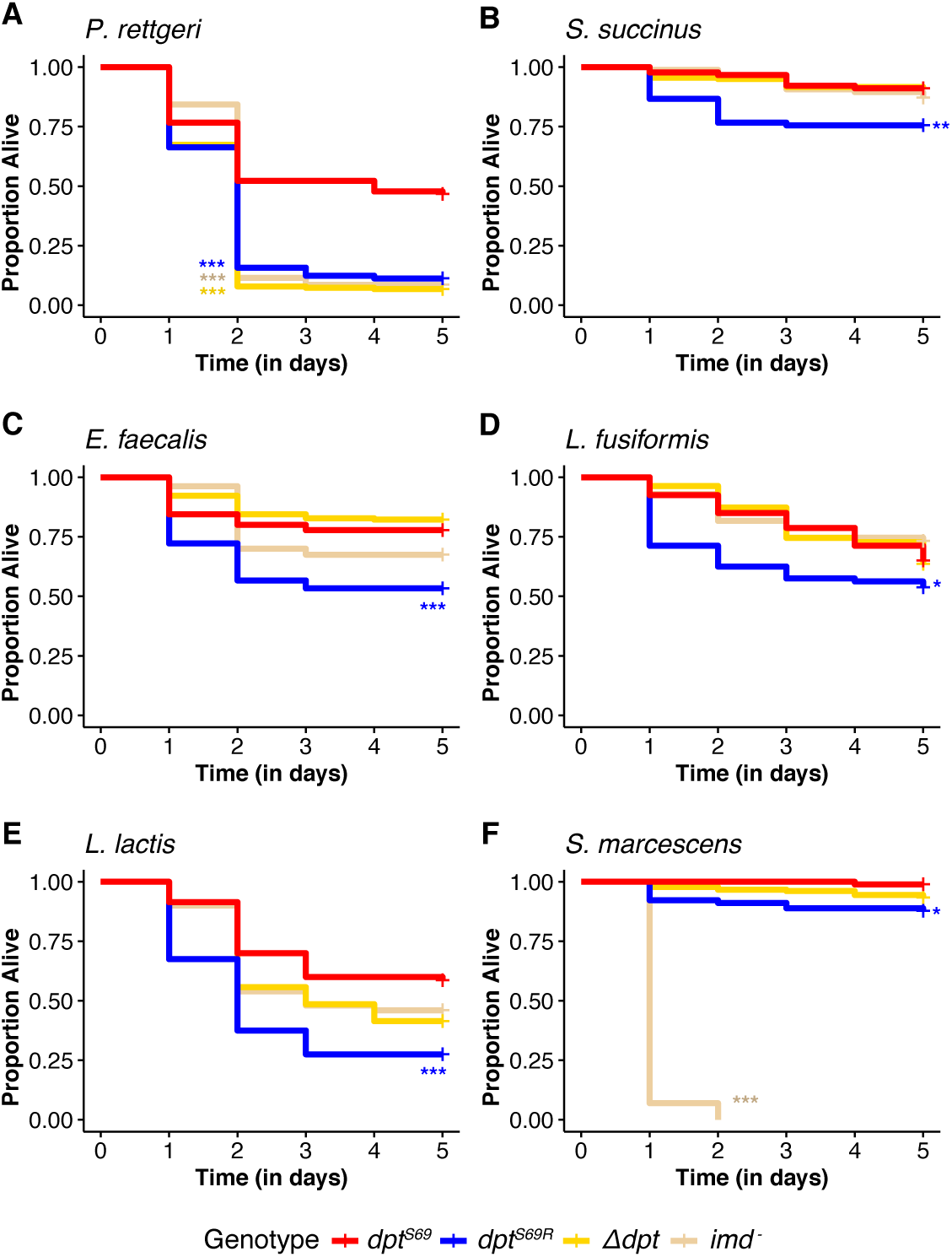
Five-day survival post systemic infection. Survival after systemic immunity challenge with the following bacteria A) *P. rettgeri,* B) *Staphylococcus succinus*, C) *Enterococcus faecalis*, D) *Lysinibacillus fusiformis* Strain Juneja, E) *Lactococcus lactis* F) *Serratia marcescens*. Each graph represents the combined results of 3 different infection dates of at least 20 males of each genotype for each date (at least 60 total). Significance is relative to dpt^S69^ (red line) using Cox proportional-hazards regression model. *P<0.05, **P<0.01, ***P<0.001.

We next challenged the CRISPR/Cas9 edited flies with systemic infection using multiple other Gram-positive and Gram-negative bacteria to determine whether the arginine allele (*dpt^S69R^*) protects against some infections better than the serine allele. Such a finding would support the hypothesis that allelic variation is maintained by different alleles providing specific protection against different microbes.

However, the differential response to systemic infection with *P. rettgeri* at an OD_600_=0.1, as described above, remains the largest difference in immune response between *Dpt* genotypes. *Dpt^S69^*flies survived better than *dpt^S69R^* flies for all systemic infections tested (Fig 1, Table S2). In the Gram-positive bacterial infections (*Enterococcus faecalis*, *S. succinus, L. fusiformis,* and *L. lactis)*, *dpt^S69R^*flies had lower survival than that of *Δdpt* or *imd^-^* (null allele for the Imd gene) flies (Fig 1B-E). The only systemic infection where *dpt* genotype did not seem to matter was *S. marcescens,* where all flies survive infection well except for *imd^-^*, which die very quickly (Fig 1F).

We also tested if males and females showed differences in systemic immunity based on genotype for *P. rettgeri, E. faecalis* and *L. plantarum* infections (Fig S2, Table S3). Generally, the *Dpt* genotypes have qualitatively similar survival patterns between males and females when infected with *P. rettgeri.* Females showed lower survival compared to males after systemic infection with *E. faecalis* (p=0.0003). Higher male survival post infection was observed previously for both *E. faecalis* and *P. rettgeri* with differences ascribed to the Toll pathway [50].

Overall, flies with the *dpt^S69^* allele are better equipped to survive a systemic bacterial infection than *dpt^S69R^* flies, though this is most pronounced for *P. rettgeri*. Of the 6 bacteria tested, there was no case where *dpt^S69R^* flies survived the infection better than *dpt^S69^* flies. These results do not support the hypothesis that alleles are maintained to better combat different pathogens.

### Diptericin genotype affects lifespan of mono- and poly-associated gnotobiotically reared flies

Although our survey of systemic infections was not exhaustive, we did not find any instances where *dpt^S69R^* flies were better able to fight infection, so we turned our attention to the role of *Dpt* in gut microbiome maintenance and immunity. The gut microbiome influences several life history traits in *Drosophila* and other organisms [51–54]. To dissect how *diptericin* genotype influences microbiome maintenance, we manipulated the microbiota in CRISPR/Cas9 flies, and measured longevity and bacterial load. We began with the longevity of axenically reared flies, since it represents a baseline survival without the presence of microbes. Flies with functional copies of *diptericin* (*dpt^S69^* or *dpt^S69R^*) did not differ in lifespan for either sex (p_genotype_=0.3762), and *Δdpt* lines show a similar lifespan. However, there was a difference in overall lifespan between the sexes among the CRISPR genome edited lines (p_sex_=1.04e-6). In *dpt^S69^* or *Δdpt* female flies have a longer lifespan than male flies, as observed generally for *D. melanogaster* previously [55]. In contrast, *imd^-^* male flies have a much longer lifespan than any of the other lines tested (Fig 2A – axenic row, Table S3).

**Figure 2:**
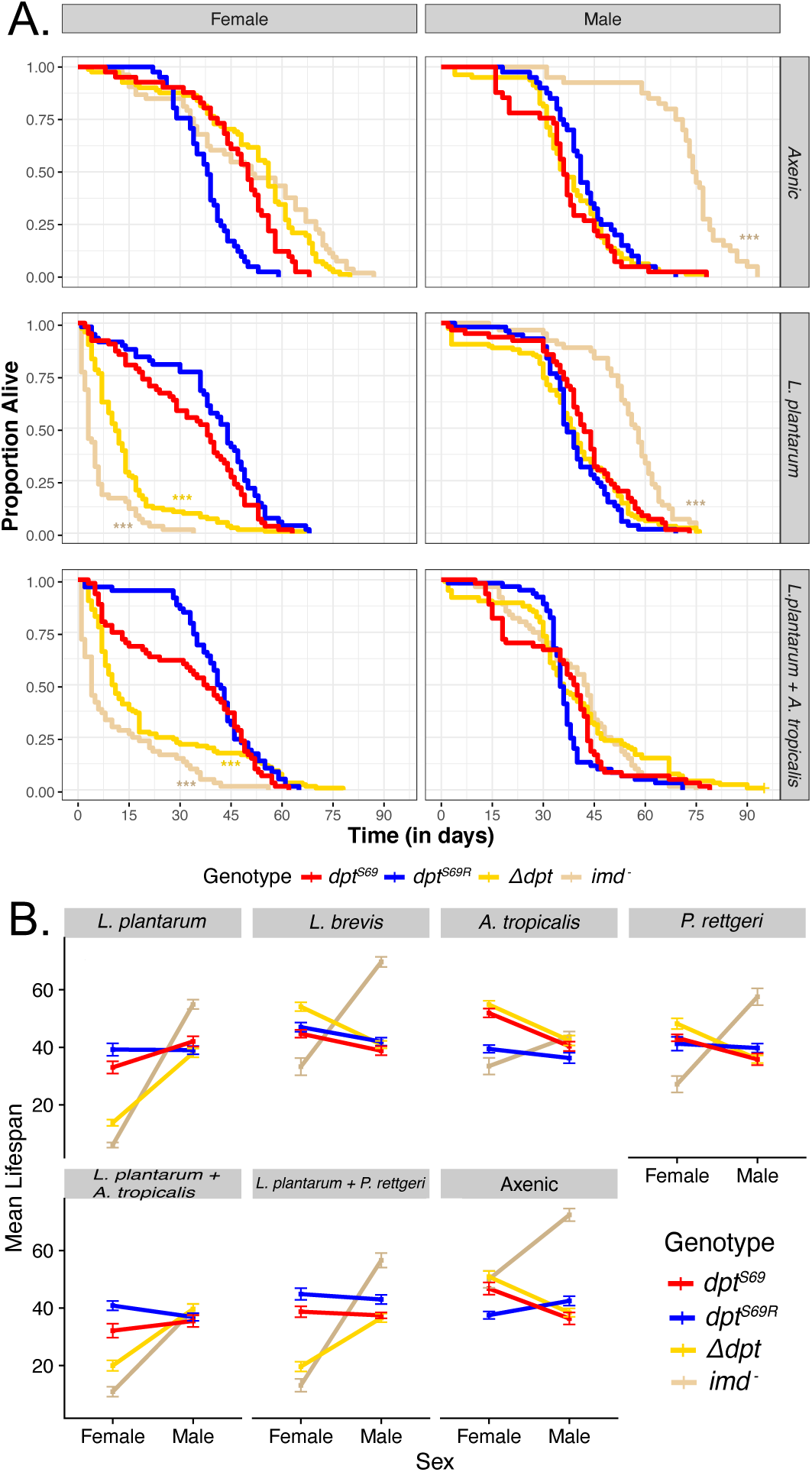
Axenic and gnotobiotic lifespan. A) Survival plots for axenic flies and gnotobiotic flies mono-associated with L. plantarum or poly-associated with L. plantarum and A. tropicalis. Each curve represents 60 flies from 3 replicates of 20 flies. Significance is shown in comparison to SS (red line) using the model Lifespan∼(Genotype*Sex)/Vial+Block. B) Mean lifespan was plotted against each sex and separated by bacterial association. Axenic flies have no bacterial association. Each point for each sex in a combination of 3 separate trials of 20 flies for a total of 60 flies. These interaction plots only have mean lifespan and thus are useful for seeing all the data at a glance. Error bars are mean plus or minus the standard error of the mean. *P<0.05, **P<0.01, ***P<0.001.

We next tested the influence of Diptericin genotype on mono-associations with standard constituents of the *Drosophila* gut microbiome. We found multiple sex effects when axenically reared flies were fed *L. plantarum, L. brevis,* or *A. tropicalis* (Figs 2A and 2B, S3). Flies that were fed *L. plantarum* show the most striking differences. Male flies continued to show similar lifespans to each other, but female *Δdpt* and female *imd^-^* flies both succumbed quickly post-feeding, indicating functional *Diptericin* is important for gut immunity against opportunistic *L. plantarum* in females.

We also fed axenic flies *P. rettgeri,* which does not readily cause gut infection, to axenically reared flies. Recall that after systemic infection *dpt^S69^* flies are more resistant to *P. rettgeri* than *dpt^S69R^* flies (Fig 1A). After monoassociation, however, we see no significant difference between the two genotypes with *P. rettgeri* in either sex (p=0.612, Fig 2B). There is also no significant difference between the *Δdpt* and lines with functional Diptericin (Fig 2B, S3). Surprisingly, however, the null allele for *Imd* again shows a significant effect on survival in a sex-specific manner: null *Imd* females die much earlier and null *Imd* males survive much longer when fed *P. rettgeri*.

Some of the sex-specific differences in survival after monoassociation with bacteria may be driven by intrinsic differences in feeding rates between the sexes (and potentially genotypes as well) [56]. To determine whether the differences in male and female survival and load were due in part to differences in feeding rates, we performed a feeding rate assay with blue dye mixed with media (Luria-Bertani broth) or *P. rettgeri* (OD_600_=15.0). We noted that females did eat more in a single hour of feeding, but that there was also an effect of *diptericin* genotype (with null flies eating less, Figure S4). Thus, it is possible that differences in longevity after monoassociation are due to different rates of exposure to those bacteria because of different feeding rates.

We next looked at how poly-associations with bacteria affected lifespan. First, we fed flies a 1:1 mixture of *L. plantarum* and *A. tropicalis*, two common gut microbes found in lab-reared and wild caught flies [57,58]. We found that female *Δdpt* and *imd^-^*flies had a much shorter lifespan than flies with functional Diptericin (Fig 2A and 2B). This is the same pattern observed in mono-association with *L. plantarum*. However, we observe that *diptericin* genotype influences survival when poly-associated with *L. plantarum* and *A. tropicalis*. *Dpt^S69R^* female flies live longer than *dpt^S69^* female flies (Fig 2A and 2B; p=0.00782). This is the opposite of systemic infections, where *dpt^S69^*flies always survived better than *dpt^S69R^* flies, and may indicate a tradeoff between defense against systemic and gut immunity.

In poly-association with 1:1 *L. plantarum* and *P. rettgeri,* we observe many of the same patterns as in the poly-association with *L. plantarum* and *A. tropicalis*. Again, female *Δdpt* and *imd^-^* flies have a shorter lifespan than female *dpt^S69^* and *dpt^S69R^* flies (Fig 2B), and *dpt* genotype is associated with lifespan. *Dpt^S69R^* females have a longer lifespan than *dpt^S69^* flies (p=0.000198, Fig 2B).

Given the genotypic and sex effects on survival after oral association with *L.* plantarum, we also looked at sex differences after systemically infecting conventionally reared flies with *L. plantarum*. We saw no difference in survival between the sexes, as observed for systemic infections with *E. faecalis* or *P. rettgeri* and saw no difference between *Δdpt* lines and lines with functional Diptericin, as observed for axenic flies mono-associated with *L. plantarum* (Fig S2). This could indicate that Diptericin plays different roles for systemic and gut immunity in relation to *L. plantarum* in each sex. Further, *dpt^S69^* male flies survived better than *dpt^S69R^* male flies (p=0.00438), in line with observations from other systemic infections in males (Fig 1).

Overall, we found a role of both specific *diptericin* genotype (serine vs. arginine) and the presence of functional copies of *diptericin* for survival after introducing common gut microbes in controlled conditions. Most striking was the sexually dimorphic role of both *Dpt* and *Imd-,* with females being much more sensitive to genotype than males.

### Gnotobiotic fly bacterial load

Given the differences in survival among sexes and genotype for different gnotobiotic associations with bacteria, we determined whether bacterial load after associations also were different among sexes and genotypes. To assess the influence of genotype on the immune response of aging flies, we studied how well common gut bacteria colonized the gut over 20 days, representing ∼20-33% of *D. melanogaster’s* normal lab lifespan [59,60]. We generated gnotobiotic flies by feeding specific bacteria for two days and observed the bacterial load 2-, 10-and 20-days post feeding.

When specifically looking at mono-association with *L. plantarum* we observe that bacterial load differences between genotypes occur within the first 2 days post feeding but begin to disappear by day 20 (p_genotype(day2)_=2.6e-5, p_genotype(day20)_=0.709, Fig 3A, S5). Note that in the first two days, the flies were raised on microbe-contaminated media, but after 2 days were moved onto sterile food and then transferred to new sterile food every 3 days. This corroborates what we saw in the longevity data in females. Within the first 15 days a large proportion of *Δdpt* female flies died, and we observed a higher bacterial load in these flies, especially on day 2 post feeding (Fig 2). By day 20, differences in bacterial load in females disappeared. Note that there is inherent sampling bias, as only the flies able to survive until day 20 are sampled at that time point. In the case of *imd^-^* flies, no females survived until day 20, hence there is no data for *imd^-^*flies on day 20 (Fig 3A, S5).

**Figure 3:**
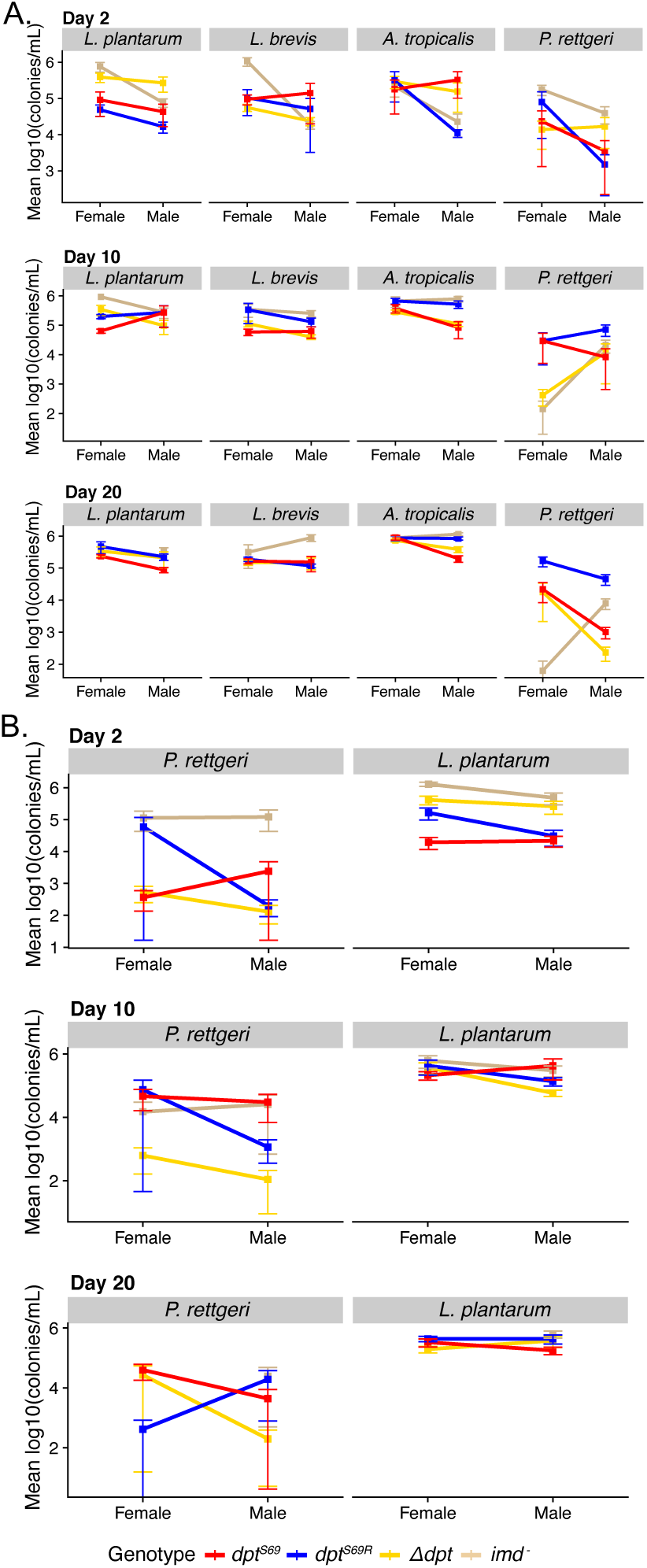
Bacterial load interaction plots for day 2, 10, and 20 post association with various microbes. Two-day old flies were placed on food seeded with a bacterial suspension of each bacteria at an OD600 of 15. Each point for each sex is from 3 separate trials of 2 samples of 3 flies each for a total of 6 samples per point. A) monoassociations where each plot represents a different condition, B) polyassociations where each row plots both the *P. rettgeri* and *L. plantarum* load from the same flies but were plated on LB and MRS respectively. Error bars are mean plus or minus the standard error of the mean.

*P. rettgeri* is the only canonical systemic pathogen we included in our gnotobiotic study, and the patterns based on genotype were different between systemic infection and the gnotobiotic oral infection data. When flies were fed *P. rettgeri,* we saw a range of bacterial loads, from zero colonies to bacterial load levels on par with common gut bacteria. We also observed a genotype effect between *dpt^S69^* and *dpt^S69R^* on day 10- and 20-post feeding in males (p_genotype(day10)_=0.0044, p_genotype(day20)_=0.0004, model only included *dpt^S69^* and *dpt^S69R^*, Table S5). In both instances, bacterial load in *dpt^S69R^* flies is higher than in *dpt^S69^* flies. This may indicate that *dpt^S69^*flies are better equipped to deal with both systemic and oral infection from *P. rettgeri*. Whether this is due to a greater ability of *dpt^S69^*flies to withstand the effects of infection (tolerance) by *P. rettgeri* by remains a question. It is also important to note that *P. rettgeri* establishes poorly in the gut of wildtype flies (personal observation), which may explain the noisy results for mono associations after oral infection.

A range of bacterial loads were also observed when flies were poly-associated with *L. plantarum* and *P. rettgeri* (Fig 3B, S6). There were no statistically significant differences between *dpt^S69^* and *dpt^S69R^* bacterial for this poly association (Table S6). In fact, there were no significant differences between any of the *P. rettgeri* bacterial loads, however *L. plantarum* does show differences on day 2-post feeding (p=7.97e-6). These differences are mainly between flies with non-functional *diptericin* (*imd-* and *Δdpt*) and flies with functional *diptericin*. Both *Δdpt* and *imd-* flies had higher bacterial load than *dpt^S69^* and *dpt^S69R^* flies on day 2 post feeding which may be an indication of the reason both these lines quickly succumb to feeding with *L. plantarum* at least in the context of the poly-association with *P. rettgeri*.

We observed larger differences in bacterial load shortly after feeding with bacteria and those differences became less by day 20-post feeding. We did not see any differences in *P. rettgeri* load when mono-associated or when part of a poly-association, indicating the flies respond differently to the same pathogen when introduced systemically or orally.

### Evidence for life history tradeoffs mediated by *diptericin* genotype

Proteins involved in a robust immune response may have pleiotropic effects on other traits either because of a direct interaction with that trait or because of inherent costs of immune defense (self-damage, energy expenditure, etc.) [61–63]. We examined three life history traits (desiccation stress survival, starvation stress survival, and uninfected longevity) to determine whether *Dpt* genotype had such pleiotropic effects.

The ability to survive desiccation stress is an important life history trait for wild *Drosophila* survival [64,65]. When subjecting our male CRISPR/Cas9 flies to desiccation, we observe conventionally reared flies succumb faster to desiccation stress than axenically reared flies (Fig 4A, p_genotype_< 2e-16). In conventionally reared males, *dpt^S69^* flies have similar desiccation resistance to *dpt^S69R^* flies (p=0.917). However, *dpt^S69^* flies survive desiccation stress better than *dpt^S69R^* flies when reared axenically (p_genotype_=0.04126). We also compared *Drosophila* OreR and W1118 (both homozygous for the serine allele of Diptericin) to the *dpt^S69^* line (Fig S7, Table S7). Unsurprisingly, despite the same *diptericin* genotype, all 3 lines show dramatically different desiccation resistance. Therefore, *diptericin* genotype plays limited, if any role in variation in desiccation resistance, as all 3 wildtype lines were derived from different genetic backgrounds.

**Figure 4:**
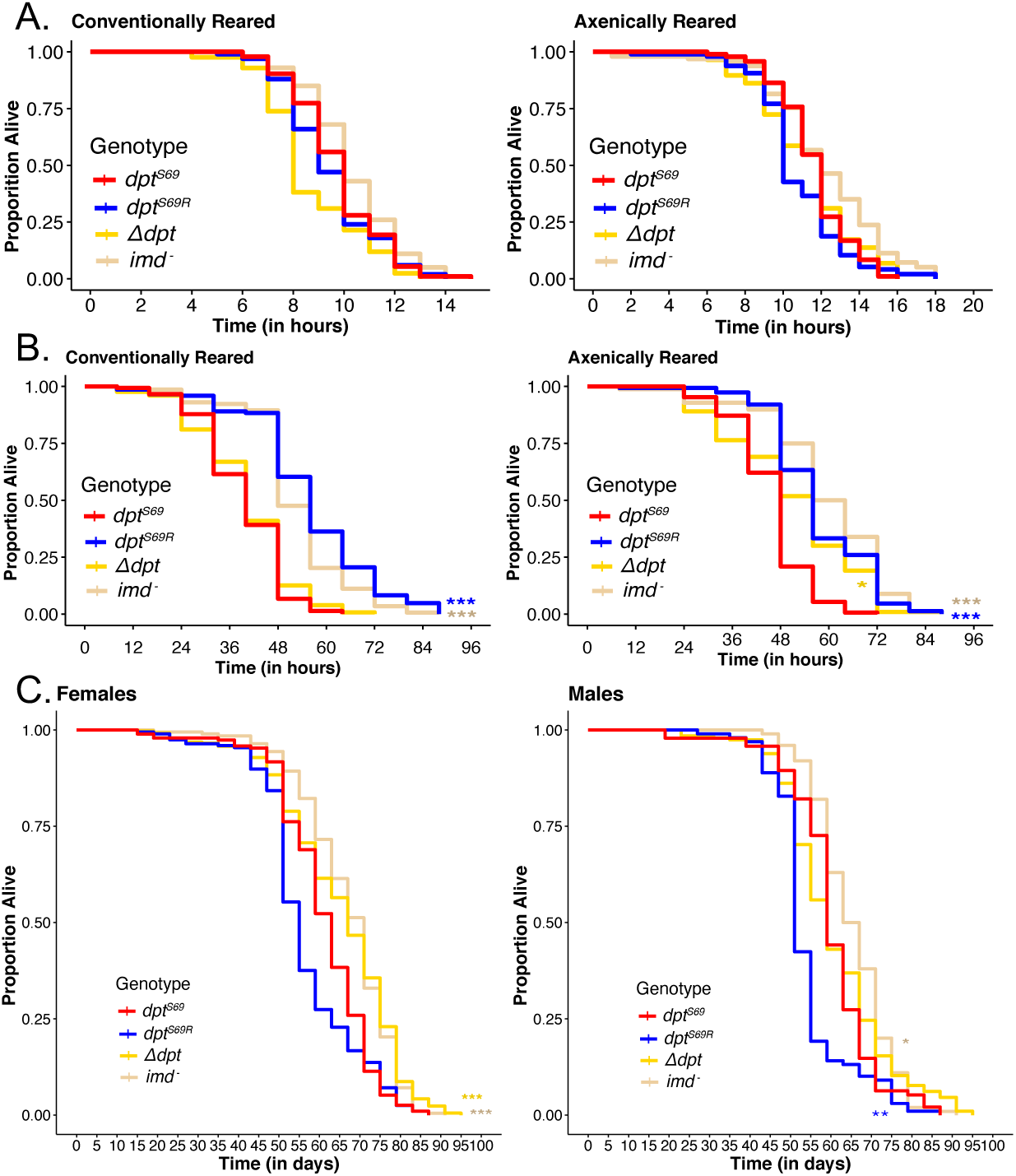
*Dpt* genotype is associated with variation in some life history traits. A) Conventionally and axenically reared male flies desiccation resistance (N>90 for each line apart from *Δdpt* line). There is no significance between *dpt^S69^* (red line) and other genotypes. B) Conventionally and axenically reared male flies starvation resistance (N>90 for each line apart from *Δdpt* line). C) Female and male lifespan for conventionally reared flies. N>93 for each line and sex. Significance is shown in relation to *dpt^S69^* (red line). Note that axenic longevity is shown in Figure 2. *P<0.05, **P<0.01, ***P<0.001.

We next looked at the effect of genetic variation in *diptericin* on male ability to survive under starvation stress. As with desiccation resistance, we found axenically reared flies survive starvation stress better than conventionally reared flies (Fig 4B, S8, p_treatment_ < 2e-16). However, unlike the desiccation stress conditions, both conventionally reared and axenically reared *dpt^S69R^*flies survive starvation stress longer than *dpt^S69^* flies (conventional: p_genotype_ < 2e-16, axenic: p_genotype_ < 2e-16). In conventionally reared flies there is no difference in survival between *dpt^S69^* flies and *Δdpt* flies (Fig 4B, Table S8). However, in axenically reared flies, *Δdpt* flies have an intermediate survival phenotype between *dpt^S69^* and *dpt^S69R^* flies. This may suggest an interaction between functional *diptericin* and the microbiome that influences starvation.

Finally, we looked at overall longevity of female and male conventionally reared flies (in the absence of any infection or other significant selection pressure). Female flies have a longer lifespan than males (Fig 4C, S9, p_sex_= 5.80e-13) regardless of *dpt* genotype. In males, *dpt^S69^* flies had a significantly longer lifespan than male *dpt^S69R^* flies (mean of 60.3 days for *dpt^S69^* and 54.1 for *dpt^S69R^*, p= 0.0072), but not in female flies (mean of 61.9 days for *dpt^S69^* and 59.0 for *dpt^S69R^,* p= 0.6434). However, in axenically reared flies only female *dpt^S69^* flies have a longer lifespan than male *dpt^S69R^* flies. Interestingly, the female flies with the longest lifespan have non-functional Diptericin (Fig 4C, *Δdpt* line and *imd^-^*line, Table S9). Males lacking functional Diptericin show the same effect, but to a lesser extent. These results provide evidence that, in the presence of a standard gut microbiota, both *Dpt* and a functional Imd pathway may decrease longevity. This is consistent with others who found downregulation of NF-κB pathways and AMPs increased lifespan in *Drosophila* [63,66], but the fact that flies with *Dpt* null alleles alone are sufficient to increase lifespan is noteworthy.

### Diptericin’s influence on gut microbial diversity

To determine whether *Diptericin* genotype influences the composition of the bacterial community in the gut, we sequenced amplicons of 16s ribosomal rRNA in conventionally reared lab flies under two different rearing conditions: flies reared in standard *Drosophila* vials, and the progeny of the cross between *dpt^S69^* and *dpt^S69R^* flies reared in cages for more than 2 generations. The vials may capture the long-term impact of genotype on microbiota but makes ruling out stochastic changes in communities difficult. The cages ensure identical microbiota during development, but do not allow gradual effects of genotype on microbiota to accumulate. First, we found flies that were co-reared in cages had similar microbiomes, with no discernable differences by the alpha diversity metric, Shannon diversity (p=0.5239, Fig 5A). On the other hand, the microbiomes of flies reared in vials was distinctly different overall compared to the microbiomes of flies reared in the cages (p<0.001, Fig 5B). However, the differences between genotypes were still minimal in the vial-reared flies and vial may be a large factor in differences between lines. There were 2 *dpt^S69R^* lines used and even though these are the same genotype and genetic background there was a significant difference between the 2 lines (Fig 5B). This difference is likely due to the within-vial drift in gut microbiome communities [67], which we attempted to control for using flies of different genotypes reared in the same cage (Fig 5A).

**Figure 5.**
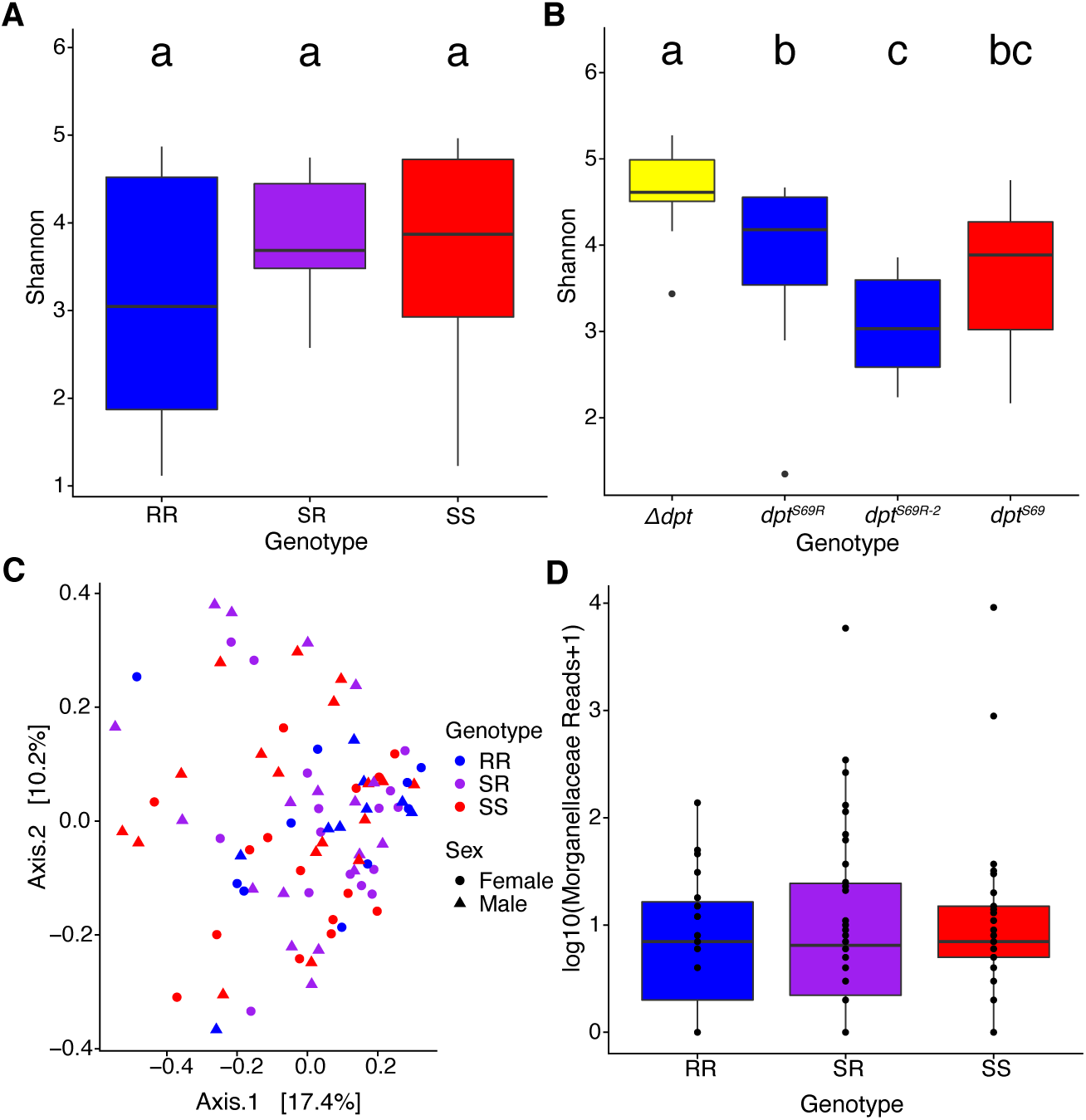
The influence of *Dpt* genotype on the gut microbiota. A) Shannon diversity of flies co-reared in cages. B) Shannon diversity of individual genotypes reared in vials. C) Bray-Curtis dissimilarity of wild caught flies. D) Differential abundance of Morganellaceae in wild caught flies. Different letters show significance based on post-hoc Tukey test, P<0.05.

The lab is a controlled environment that allows for excellent control of variables but does not replicate the conditions found in nature. Thus, we looked at the microbiomes of wild caught flies collected from the decaying fruit of an apple orchard. We found flies with homozygous for *dpt^S69R^*, homozygous for *dpt^S69^*, and heterozygous genotypes at our collection location (Table S10). Out of the 955 *D. melanogaster* flies successfully genotyped, only 20 flies were homozygous for *dpt^S69R^*were identified. All 20 homozygous *dpt^S69R^* flies were profiled along with 36 each of *dpt^S69^* homozygous flies and heterozygous flies for a total of 92 flies profiled (amplicon sequences of both *Dpt* and 16s rRNA are in Table S2).

We found that *dpt* genotype does not affect the overall composition of the microbiome of wild caught flies based on Shannon diversity (alpha diversity) and Bray-Curtis dissimilarity (beta diversity) (Fig 5C). However, when looking at differential abundance of individual bacterial families there are differences based on genotype. We tested specific association between *diptericin* genotype and normalized counts from 9 microbial families. Most intriguing is the difference in abundance of *Morganellaceae* family reads since, *P. rettgeri* belongs to this family. Heterozygous and homozygous serine flies have a slightly higher abundance than homozygous arginine flies (Fig 5D), but these differences were not significant.

Overall, our results for associations between *Dpt* genotype and gut microbiome in both lab and wild flies are weak. Such associations may require much larger sample sizes if the effects are small – particularly given the noisy phenotype in the wild. Alternatively, the influence of *Dpt* genotype on gut microbiome diversity may be specific to developmental life stages or microbes that are relatively rare in these populations.

## Discussion

The adaptive maintenance of multiple alleles of a single gene has been posited for nearly a century [68–72] and evidence for its pervasiveness continues to grow [73–78]. However, there are relatively few cases where we understand the adaptive benefits of both allele (sickle-cell anemia and malaria resistance being the best studied example [79]). In this study, we generated CRISPR/Cas9 genome-edited flies of different *diptericin* genotypes to investigate mechanisms of maintenance of allelic variation in AMPs. This allowed us to specifically study a single amino acid change on an otherwise genetically-controlled background. Overall, we found evidence that *dpt^S69^* flies survive systemic infection better, while *dpt^S69R^*flies survive some opportunistic gut infections better. Thus, more robust defense against systemic infection appears to interfere with maintaining a balanced gut microbiota – this tradeoff may result in natural populations maintaining both alleles. Importantly, we note that while our results are consistent with the adaptive maintenance of alleles in natural populations, these lab assays do not prove the adaptive value of different alleles in the field. Systemic infections with *P. rettgeri* in CRISPR/Cas9 genome-edited lines confirmed the amino acid polymorphism in Diptericin as the basis for the difference in immune response of flies with different *diptericin* genotypes previously hypothesized *via* an association study [19].

Surprisingly, *dpt^S69^* flies had a higher survival five days post infection than *dpt^S69R^* flies in *all* systemic infections tested. This may mean either that the serine allele provides improved defense against all the tested systemic infections, or that alterations to other aspects of fly physiology (notably the gut microbiota) could predispose arginine flies to poorer survival. It is also possible that some of *diptericin’s* influence on survival is mediated by changes in the microbiome. We did not perform systemic infections on axenically-reared flies, but if *Dpt* genotype influences the microbiome, which in turn influences survival after infection with other pathogens, *Dpt* should not influence survival after systemic infection if flies are reared axenically.

Our study also reveals an important role of Diptericin in preventing opportunistic gut infections by common gut microbes. This is especially evident when looking at lifespan after associating axenic flies with *L. plantarum* in both mono- and poly-association contexts. In both sexes, *dpt^S69R^* flies have a longer lifespan than *dpt^S69^* flies, and this effect is enhanced in poly-association with *L. plantarum* and *A. tropicalis.* Other studies have posited a role for individual AMPs in the maintenance of gut microbes in *Drosophila* [29], but this appears to be the first example linking a functional copy of a single AMP to survival differences.

One of our most striking and unexpected results was the distinct sex differences in longevity after association with common gut bacteria. Furthermore, fly genotype plays a significant role in these differences. After association with *L. plantarum*, *imd-* females had the shortest mean survival, while *imd-* males had the longest mean survival. The effect for *Δdpt* was similar: females had much shorter average survival than the other two *dpt* alleles, but the three alleles showed equivalent longevity in males. While the *imd-* sexual dimorphism is relatively consistent across experiments with different microbes, the *Δdpt* effect seems to be more limited to associations involving *L. plantarum.* This result not only highlights the importance of using both sexes in microbiome research, but also confirms that functional Diptericin is important for *Drosophila melanogaster* gut health. There is a growing body of literature on the effects of sexual dimorphisms in immune response, but it is lacking for sexual dimorphisms in gut immunity and our findings only emphasize the need to fill this gap (*Drosophila* immunity sexual dimorphisms reviewed in Belmonte et al. 2020; also see [80]). Our feeding assay provides some evidence that the differences between male and female survival after exposure to microbes is due the amount of food (and therefore the number of microbes) consumed, but these differences should be examined in much more detail. More generally, the sex by genotype effect suggests a likely explanation for the maintenance of genetic variation: different fitness effects of alleles in the different sexes [81]. Note that while our study focuses on the serine/arginine polymorphism, at least 6 null alleles of *dpt* segregate in natural populations of *D. melanogaster* and the frequency of those nulls shows clinal variation in both North American and Africa [82], thus the nulls are likely maintained selectively too.

Finally, we note that we have framed our argument in terms of balancing selection in the broad sense because we are agnostic as to the population dynamics upon which these different selective pressures might act. For example, it may be that some populations are disproportionately infected by *P. rettgeri* or other microbes for which the serine allele is favored, while other populations are relatively pathogen-free and the arginine allele is favored. This scenario is partly supported by the evidence of clinal patterns in allele frequency in Africa and (to a lesser extent) North America [83]. Similarly, these selective pressures could vary temporally. On the other hand, these pressures could act all in one population where serine is favored in infected individuals and the arginine allele is favored in uninfected individuals.

Herein we highlight 3 main roles of Diptericin in *Drosophila*. 1) Diptericin genotype influences systemic immune defense; 2) in certain conditions Diptericin genotype influences gut immunity; and 3) Diptericin has sex-specific effects in the gut. These results highlight the need for individuals and populations to modulate the immune system to balance systemic and gut immunity, and how the different needs of females and males complicate this balance. The dramatic differences in survival of males and females in response to oral infection by common gut bacteria underlines the importance of looking at both sexes when examining the maintenance of genetic diversity, as balancing selection may be caused by sexual dimorphism. Overall, our results suggest that a complex interaction between sex, environmental context (starvation, pathogen exposure – both systemic and oral), and genotype may contribute to the long-term maintenance of immune alleles.

## Materials and Methods

### Drosophila Lines and Rearing

Conventionally reared flies were maintained in a 23°C incubator with a 12h light:12h dark schedule on a cornmeal-molasses-yeast diet (64.3g/L cornmeal, 79.7mL/L molasses, 35.9g/L yeast, 8g/L agar, 15.4mL of food acid mix (50mL Phosphoric Acid + 418mL Propionic Acid + 532mL deionized water) and 1g/L Tegosept. We used CRISPR/Cas9 genome editing to modify the *diptericin A* gene. Briefly, DNA coding for guide RNA (gRNA) was inserted into the pUS-BbsI plasmid (Table S1). A single stranded donor DNA (120bp) containing the desired edit (to change from serine to arginine) and a silent mutated PAM site was synthesized by IDT (Coralville, IA, USA). The plasmid and ssODN were then injected into Bloomington stock #55821, which expresses Cas9 driven by the *vasa* promoter, by Genetivision, Inc. (Houston, TX, USA). Individual flies developed from the injected embryos were collected and crossed with a modified version of Bloomington stock # 7198 (a line with *CyO*/*Kruppel* balanced on the 2^nd^ chromosome, and *serrate*/*Dichaete* balanced on the 3^rd^ chromosome). Our version, 7198^A4^, was provided by Stuart Macdonald and has the DSPR [84] A4 line’s X chromosome instead of the *w*[*] from the original 7198. The F1s were collected and again individually crossed with 7198^A4^ for F2 crosses yielding individuals with a homozygous 2^nd^ chromosome representing one of the chromosomes carried by the original injected embryo. The Dpt gene was sequenced from the F2 cross progeny to determine whether edits occurred. This yielded several classes of alleles including homozygous serine *dpt* (wildtype, *dpt^S69^*), homozygous arginine *dpt* (*dpt^S69R^*), and *dpt* null (*Δdpt,* refers to lines with either 1 or 3 base pair deletion) (Fig S1A). Balancers were removed, and lines were moved into the same genetic background through a series of crosses as shown in Fig S1B. An *imd-* line was used as an IMD pathway negative control but note that this line was from a completely different genetic background from the rest of the lines.

### Axenic Fly Preparation

Microbe-free (axenic) lines were generated by first washing embryos in a 10% bleach solution to dissolve the chorion for 2 minutes. The embryos were then washed in 70% ethanol for 30 seconds and water for another 30 seconds, then transferred to autoclaved molasses food (see above). Some embryos from each treatment were placed onto De Man, Rogosa and Sharpe (MRS) agar plates and incubated at 30°C for 48 hours to check that they did not contain viable microbes. Axenic lines were continuously checked for the presence of contaminating microbes (every 3-4 generations) by homogenizing flies and plating the homogenate on MRS agar.

Axenically and gnotobiotically (see below) reared flies were maintained in an incubator that was isolated from conventionally reared flies. The incubator was kept at 23°C with a 12h light:12h dark schedule. Axenic and gnotobiotic flies were kept on the same molasses diet that had been autoclaved before dispensing into autoclaved vials. Axenic and gnotobiotic flies were only handled inside a sterile hood (Baker SG 400, The Baker Company Inc., Sanford, ME).

### Bacterial Strains

The following bacteria were used for systemic infection assays: *Providencia rettgeri* [85]*, Providencia burhodegraneria* Strain B [85]*, Enterococcus faecalis* [85])*, Serratia marcescens* [40]*, Lysinibacillus fusiformis* Strain Juneja [86], and *Staphylococcus succinus* (isolated from wild *Drosophila,* Unckless lab). Bacteria were grown from glycerol stocks on LB plates at 37°C overnight.

The following bacteria were used in gnotobiotic experiments: *L. plantarum, L. brevis, A. tropicalis.* All these strains were isolated from plating conventionally reared flies on MRS agar in the Unckless lab at the University of Kansas. Individual colonies from plated fly homogenate were grown overnight in MRS for DNA isolation. Bacterial species were identified using Sanger sequencing with the 16S rRNA region primers 27F (AGAGTTTGATCCTGGCTCAG) and 1492R (CGGTTACCTTGTTACGACTT). We also utilized the same *P. rettgeri* as described for systemic infection.

### Survival Assays

For systemic infections, individual colonies of bacteria were picked and grown in 2mL LB broth shaking overnight at 37°C. Bacterial suspensions were then diluted or concentrated to OD_600_=0.1 for *P. rettgeri,* OD_600_=10 for *S. succinus,* OD_600_=1.5 for *E. faecalis*, OD_600_=3.5 for *L. fusiformis*, OD_600_=1.0 for *L. lactis*, and OD_600_=4.0 for *S. marcescens*. *L. plantarum* was grown in 5mL MRS at 30°C overnight and was concentrated to OD_600_=10 for systemic infections. To induce systemic infection, 5-9 day old, conventionally reared flies were pricked in the thorax with a needle dipped in a bacterial suspension [87]. Infections were done in triplicate with at least 20 flies for each replicate per line for a total of 60 flies per genotype per condition. Flies were incubated at 23°C with a 12h light:12h dark schedule and survival was tracked daily for 5 days post infection.

### Gnotobiotic Longevity

Axenically reared flies were collected within 24 hours of eclosion. Flies were then kept on sterile food for 2 days before sorting for longevity experiments. To begin longevity experiments, flies were separating into groups of 10 individuals of each sex and put onto sterile food seeded with 50uL of bacterial suspension at an OD600 of 15 +/-1. For each replicate, we used 2 vials of 10 flies each per sex per line for a total of 20 flies for each sex per genotype for a total of 60 flies across all replicates. Flies were allowed to feed in the inoculated vials for 3 days before being transferred to uninoculated sterile food vials. Flies were flipped to new sterile media every 4-5 days for the remainder of the experiment. Surviving flies were counted every 1-3 days until all flies were dead.

### Gnotobiotic Bacterial Load

To determine whether microbes became established in the gut, we homogenized flies during and after the exposure and plated the homogenate. We measured bacterial load by inoculating flies in the same manner as gnotobiotic longevity. Flies were separated into groups of 5 females and 5 males per vial. For 2-day experiments flies were kept on the seeded food for the entire experiment. For 10– and 20-day experiments, flies were allowed to feed on the seeded food for 3 days before being transferred to sterile food. Flies continued to be transferred to new sterile media every 3-4 days until day 10 or 20 post feeding. After the experimental (2, 10 or 20 day) time period, flies were surface sterilized by washing in 70% ethanol followed by molecular grade water. Flies were separated by sex and three individuals were homogenized together in 300uL of sterile 1x PBS and the homogenate was plated on the appropriate media using a Whitley WASP Touch® spiral plater (Don Whitley Scientific, UK). When there were not 3 flies still alive then all remaining flies of a sex were used and squished in 100uL of 1x PBS per fly collected instead of 300uL to keep all samples the same concentration per fly. Counts were adjusted accordingly.

### Feeding Rate Assays

To measure the amount of food and bacteria consumed by males and females of different genotypes, we made food containing blue dye by adding 11.2g FCF blue dye (Erioglaucine disodium salt) per liter of food. We used newly eclosed flies (1-2 days post eclosion / 14 days post oviposition) and separated sexes (in sterile conditions) and keep flies at a density of 10 flies/vial in fresh food vials. We held these flies in incubators for one day, so they would recover stress of anesthesia during sexing. To introduce bacteria (or control media) into the food, we pipetted 50uL of suspension or LB at an OD600 of 15 (+/-1) into each vial and allowed the suspension to absorbed into the food for 30-40 mins by keeping the vials open inside a sterile hood. We next added the experimental flies and allowed them to feed for one hour. After 1 hour, we anesthetized the flies on ice, then rinsed in ethanol and sterile water. Flies were homogenized in 300µl of 1X PBS with a glass bead (maximum speed for 4 minutes). The homogenate was centrifuged at 14000 RPM for 4 minutes and 200µl of the supernatant was used to measure absorbance at 630nm. Absorbance differences were analyzed using the natural log of absorbance as the response variable with genotype, sex, genotype by sex interaction, and block as independent variables. Due to the blocking structure of the experiment, each treatment (no media control, LB control, *P. rettgeri*) was analyzed separately.

### Desiccation

Desiccation survival assays were performed on *dpt^S69^*and *dpt^S69R^* adult males 4-7 days post-eclosion in conventionally and axenically reared flies. Ten males were placed into an empty vial and closed off with a cotton flug. Each genotype had 5 vials for a total of 50 flies per line per rearing condition in each replicate. The flugs were topped with silica gel (Fisher, #S684) and sealed with parafilm to prevent any moisture from entering the vials. Vials were kept at 24°C on 12 hour day/night cycles. Survival was measured by counting flies hourly until the entire population died. This was repeated once more with 50 flies per line per rearing condition in each trial for a total of 100 flies with the exception for the *Δdpt* line which in total only had 30 flies per condition in total across trials.

### Starvation

Starvation survival assays were performed on *dpt^S69^*and *dpt^S69R^* adult males 4-7 days after eclosion for axenically and conventionally reared flies. Ten males were placed into a vial with autoclaved starvation media (1% agar). The 1% agar was used to starve flies of nutrition but not desiccate them. Vials were kept at 24°C on 12-hour day/night cycles. Survival was measured by counting surviving flies at three 8-hour intervals (8 am, 4 pm, and 12 am) until all flies died. This was repeated twice more for a total of 3 trials with at least 40 flies per line per rearing condition in each trial for a total of at least 120 flies.

### 16S Sequencing

Flies were reared in the lab in two distinct ways for 16S rRNA sequencing of conventionally reared flies. First, flies were taken from vials of individual genotypes. This allows for any moderate effects of fly genotype to equilibrate over time. Second, flies were taken from cages that were started with heterozygous *diptericin* flies. These cages were started with the F1 progeny from crosses of *dpt^S69^* and *dpt^S69R^* flies and allowed to continue for 3 discrete generations before flies were collected for 16S rRNA sequencing. This ensures that the genotypes are exposed to the same microbes and any differences in microbiome are due to genetic differences manifest in that generation.

We isolated DNA from individual flies using Gentra PureGene Tissue DNA Isolation Kit (Qiagen #158388, Qiagen, Germantown, MD) following manufacturer’s instructions. DNA pellets were rehydrated in 10 μl DNA hydration solution. Flies from cage populations were genotyped by using the dpt_cw primer pair (Table S1) followed by a restriction enzyme digest using enzymes that cut DNA sequences for arginine (BccI, NEB # R0704S) or serine (AluI, NEB # R0137S).

DNA concentration was measured with Qubit fluorimeter (Invitrogen). Ten flies from each inbred CRISPR genome edited line (5 females and 5 males from *dpt^S69^*, *dpt^S69R^*, *dpt^S69R-2^*(a second homozygous arginine line from CRISPR/Cas9 genome editing), and *Δdpt* (1 base pair deletion) and ten flies of each genotype from the cage population (5 females and 5 males of SS, SR, RR genotypes) were each brought to a DNA concentration of 5 ng/μl. Libraries were prepped in accordance with the 16S Metagenomic Sequencing Library Preparation protocol from Illumina for the 16S V3/V4 region. Sequencing was performed on the Illumina MiSeq platform using v3 300bp paired end reads. Library preparation and sequencing was performed at the University of Kansas Genome Sequencing Core (Lawrence, KS, USA).

Wild flies were collected from decaying apples and pears in an apple orchard in Kansas City, KS (3341 N 139^th^ St, Kansas City, KS 66109). Flies were immediately transported back to the lab and sorted by species on CO_2_ and frozen at -20°C. DNA was extracted from individual flies using Gentra PureGene Cell & Tissue DNA Isolation Kit (Qiagen #158388). The samples were tested for species (*D. simulans* vs. *D. melanogaster*)*, Wolbachia* status, and Dpt genotype using primers listed in Table S1. Collections are summarized in Table S10. Libraries were prepped in the same manner as the conventionally reared flies.

### 16S Bioinformatic Analysis

Demultiplexed reads were processed with QIIME2, v 2019.10 [88]. Primers were removed from 5’ ends with Cutadapt using default parameters [89]. Reads were de-noised and trimmed for quality with Divisive Amplicon Denoising Algorithm (DADA2) within the QIIME2 bioinformatics pipeline [90]. Forward and reverse reads were truncated at 280 bp and 245 bp respectively. The remaining ASV table was exported from QIIME2 for further processing in R [91]. Taxonomy was assigned to the ASV table using SILVA 16S rRNA gene reference database, v.138 [92–94]. Reads assigned to Genus level *Wolbachia* (a *Drosophila* endosymbiont) and Kingdom level Eukaryota were removed from further analysis. We also removed reads not observed at least 3 times in at least 10 samples. Then, conventionally reared flies were rarefied to 17066 reads, while wild flies were rarefied to 10771 reads.

All statistical analysis of 16S data was performed in R (4.1.2) using the Phyloseq [95] package and ggplot2 [96] package for visualization. The CRISPR and cage population data were analyzed separately. For each population, alpha diversity was estimated using Shannon diversity in Phyloseq using the estimate_richness() function. Bray-Curtis dissimilarity was calculated to look at overall patterns of microbiome composition in Phyloseq using the ordinate() function.

Significance was determined at α=0.05. Fixed effects models were fit with the package lme4 [97] with the fixed effects genotype + sex + genotype*sex. When necessary, P-values were adjusted for multiple comparisons using FDR correction method.

## Statistical Analysis

R (version 4.1.2) was used to run statistical analyses. Survival data were plotted using the R package survminer [98]. The analysis was performed using the Cox proportional-hazards regression model in R [99]. For longevity, significance was determined using the model: Lifespan∼(Genotype*Sex)/Vial+Block. For gnotobiotic bacterial load, significance was determined using the model: Colonies.mL∼(Genotype)*Sex+Block.

## Competing Interests

The authors state no competing interests.

## Acknowledgments

We thank the Cider Hill Family Orchard for allowing us to collect wild flies on their property; S. Macdonald lab for providing the modified 7198 and A4 lines; Kistie Brunsell for help with DNA extractions of wild flies; and Elizabeth Everman for assistance with the feeding assay. The University of Kansas Genome Sequencing Center Core (supported by NIH CMADP COBRE P20-GM103638) performed 16s sequencing. SRM was supported by a University of Kansas Self Graduate Fellowship, the work was supported by NIH R01-AI139154 to RLU.

## Supplemental tables

**Table S1.**
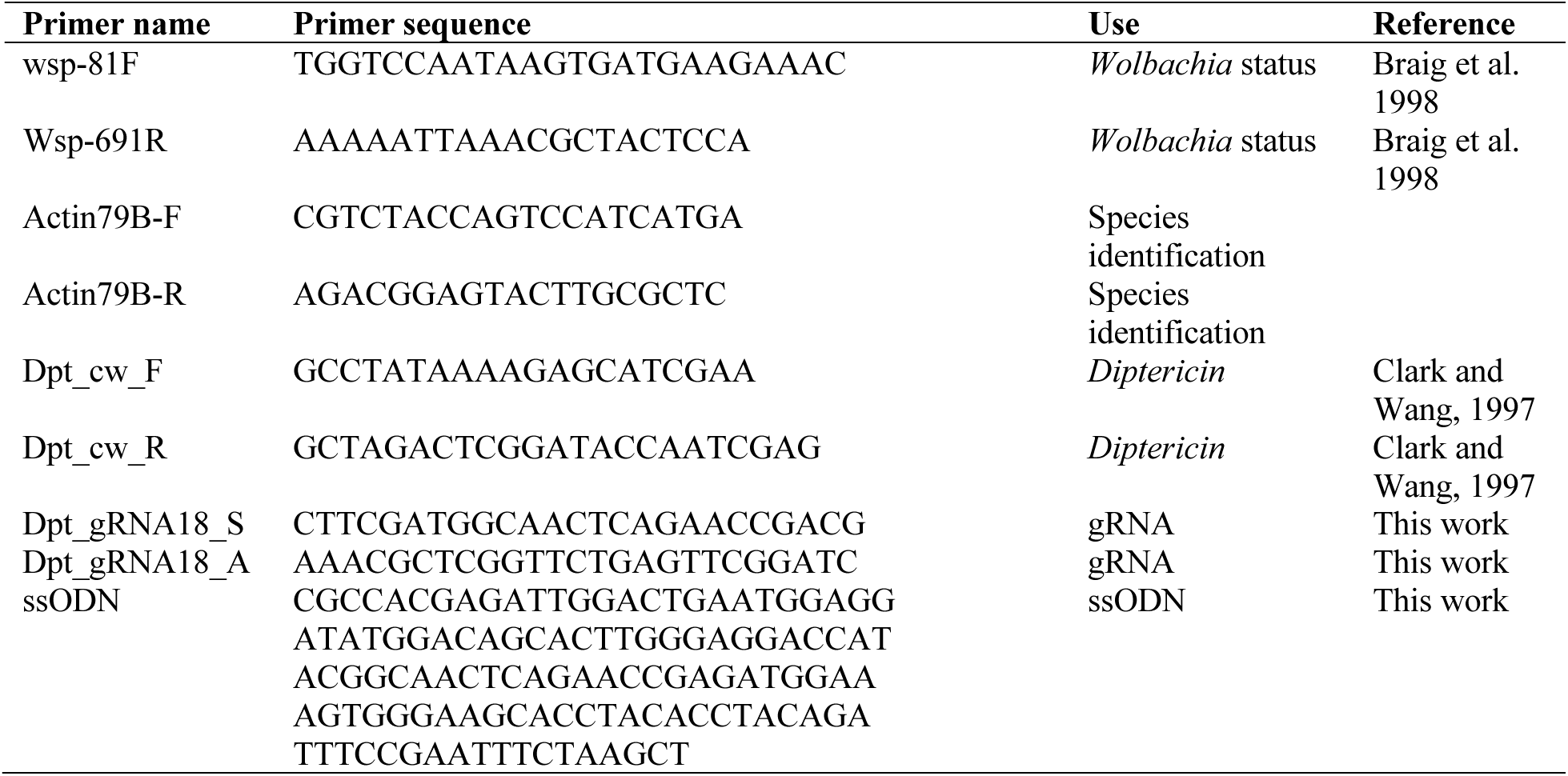
Oligonucleotide sequences used in this study.

**Table S2.** Cox proportional hazard pairwise significance p-values for systemic infection. P-values less than 0.05 are underlined.

|  |  |  |  |
| --- | --- | --- | --- |
| ***P. rettgeri*** |  |  |  |
| | *imd-* | $\Delta dpt$ | *dptS69R* |
| $\Delta dpt$ | <u>4.07E-02</u> | - | - |
| *dptS69R* | 4.54E-01 | 0.26329 | - |
| *dptS69* | <u>1.20E-04</u> | <u>1.10E-09</u> | <u>1.30E-06</u> |
| ***S. succinus*** |  |  |  |
| | *imd-* | $\Delta dpt$ | *dptS69R* |
| $\Delta dpt$ | 0.4994 | - | - |
| *dptS69R* | 0.0672 | <u>0.0023</u> | - |
| *dptS69* | 0.4994 | 0.9828 | <u>0.012</u> |
| ***E. faecalis*** |  |  |  |
| | *imd-* | $\Delta dpt$ | *dptS69R* |
| $\Delta dpt$ | <u>0.0264</u> | - | - |
| *dptS69R* | <u>0.0264</u> | <u>1.00E-06</u> | - |
| *dptS69* | 0.2969 | 0.3405 | <u>2.10E-03</u> |
| ***L. fusiformis*** |  |  |  |
| | *imd-* | $\Delta dpt$ | *dptS69R* |
| $\Delta dpt$ | 0.435 | - | - |
| *dptS69R* | <u>0.048</u> | 0.22 | - |
| *dptS69* | 0.435 | 0.927 | 0.19 |
| ***L. lactis*** |  |  |  |
| | *imd-* | $\Delta dpt$ | *dptS69R* |
| $\Delta dpt$ | 0.7515 | - | - |
| *dptS69R* | <u>0.0474</u> | <u>0.0474</u> | - |
| *dptS69* | 0.1966 | 0.0858 | <u>0.0015</u> |
| ***S. marcescens*** |  |  |  |
| | *imd-* | $\Delta dpt$ | *dptS69R* |
| $\Delta dpt$ | <u><2e-16</u> | - | - |
| *dptS69R* | <u><2e-16</u> | 0.1151 | - |
| *dptS69* | <u><2e-16</u> | 0.054 | <u>0.0042</u> |

**Table S3.**
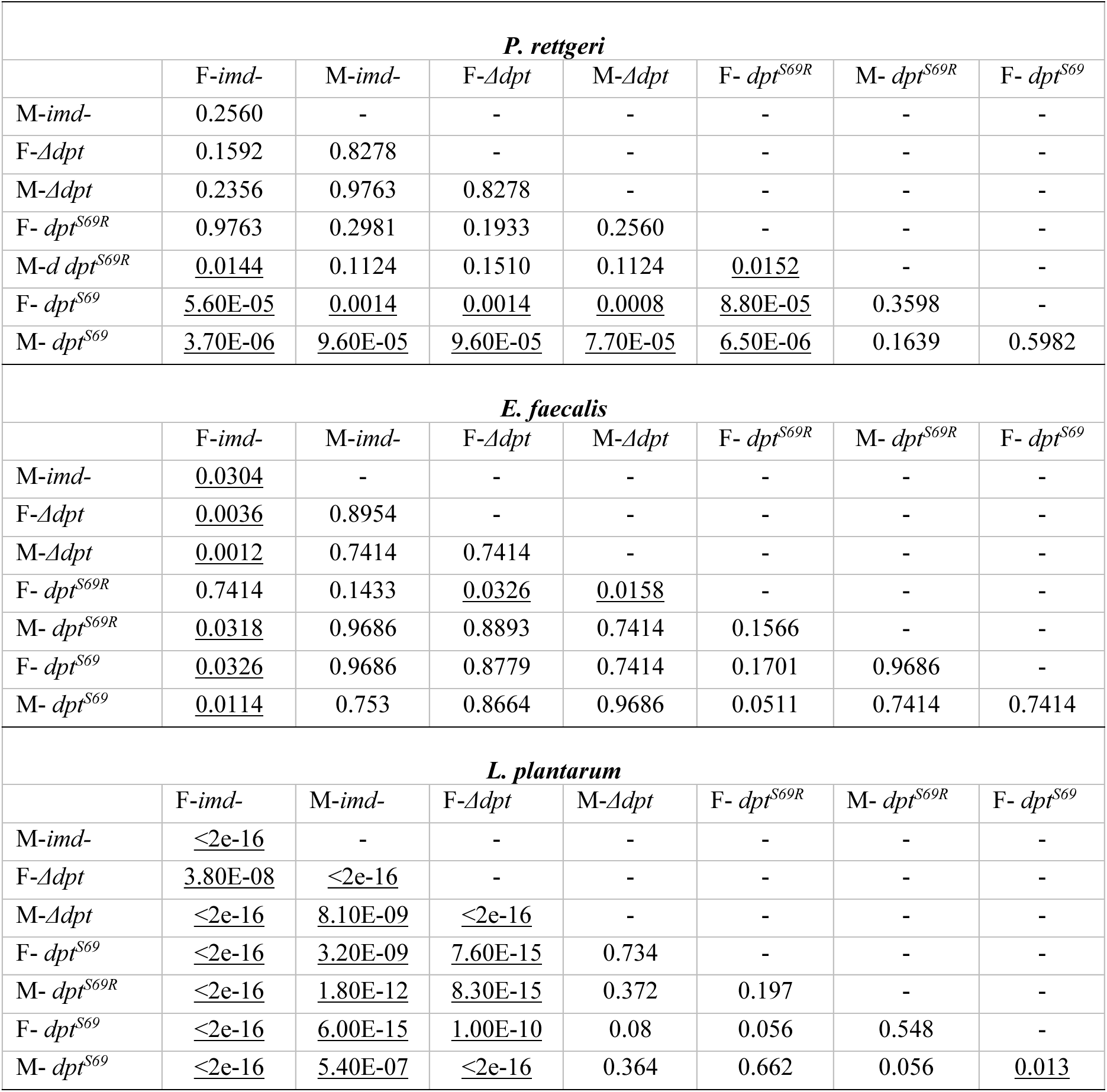
Cox proportion hazard ratio pairwise p-values for systemic infections with males and females. P-values less than 0.05 are underlined.

**Table S4.** P-values from Tukey HSD for axenic and gnotobiotic longevity. P-values less than 0.05 underlined.

| Axenic |  |  |  |  |  |  |  |
| --- | --- | --- | --- | --- | --- | --- | --- |
|  | F-imd- | F-Δdpt | F-dpt<sup>S69R</sup> | F-dpt<sup>S69</sup> | M-imd- | M-Δdpt | M-dpt<sup>S69R</sup> |
| F-Δdpt | 1 | - | - | - | - | - | - |
| F-dpt<sup>S69R</sup> | <u>6.00E-04</u> | <u><2e-16</u> | - | - | - | - | - |
| F-dpt<sup>S69</sup> | 0.9447 | 0.8051 | 0.063 | - | - | - | - |
| M-imd- | <u><2e-16</u> | <u><2e-16</u> | <u><2e-16</u> | <u><2e-16</u> | - | - | - |
| M-Δdpt | <u>1.00E-04</u> | <u><2e-16</u> | 1 | <u>0.0492</u> | <u><2e-16</u> | - | - |
| M-dpt<sup>S69R</sup> | 0.1713 | 0.0519 | 0.7467 | 0.8822 | <u><2e-16</u> | 0.8133 | - |
| M-dpt<sup>S69</sup> | <u>1.00E-04</u> | <u><2e-16</u> | 1 | 0.0218 | <u><2e-16</u> | 0.9947 | 0.5114 |
| *L. plantarum* |  |  |  |  |  |  |  |
|  | F-imd- | F-Δdpt | F-dpt<sup>S69R</sup> | F-dpt<sup>S69</sup> | M-imd- | M-Δdpt | M-dpt<sup>S69R</sup> |
| F-Δdpt | <u>0.0064</u> | - | - | - | - | - | - |
| F-dpt<sup>S69R</sup> | <u><2e-16</u> | <u><2e-16</u> | - | - | - | - | - |
| F-dpt<sup>S69</sup> | <u><2e-16</u> | <u><2e-16</u> | 0.185 | - | - | - | - |
| M-imd- | <u><2e-16</u> | <u><2e-16</u> | <u><2e-16</u> | <u><2e-16</u> | - | - | - |
| M-Δdpt | <u><2e-16</u> | <u><2e-16</u> | 0.9992 | 0.2479 | <u><2e-16</u> | - | - |
| M-dpt<sup>S69R</sup> | <u><2e-16</u> | <u><2e-16</u> | 1 | 0.2568 | <u><2e-16</u> | 0.9999 | - |
| M-dpt<sup>S69</sup> | <u><2e-16</u> | <u><2e-16</u> | 0.9531 | <u>0.0056</u> | <u><2e-16</u> | 0.5593 | 0.9198 |
| *P. rettgeri* |  |  |  |  |  |  |  |
|  | F-imd- | F-Δdpt | F-dpt<sup>S69R</sup> | F-dpt<sup>S69</sup> | M-imd- | M-Δdpt | M-dpt<sup>S69R</sup> |
| F-Δdpt | <u><2e-16</u> | - | - | - | - | - | - |
| F-dpt<sup>S69R</sup> | <u>1.00E-04</u> | 0.1564 | - | - | - | - | - |
| F-dpt<sup>S69</sup> | <u><2e-16</u> | 0.5911 | 0.9977 | - | - | - | - |
| M-imd- | <u><2e-16</u> | <u>1.12E-02</u> | <u><2e-16</u> | <u>1.00E-04</u> | - | - | - |
| M-Δdpt | <u>2.34E-02</u> | <u><2e-16</u> | 4.81E-01 | 9.95E-02 | <u><2e-16</u> | - | - |
| M-dpt<sup>S69R</sup> | <u>0.0013</u> | <u>3.63E-02</u> | 0.9997 | 0.9434 | <u><2e-16</u> | 0.8429 | - |
| M-dpt<sup>S69</sup> | 0.0938 | <u>1.00E-04</u> | 0.6415 | 0.2208 | <u><2e-16</u> | 1 | 0.9079 |
| *L. brevis* |  |  |  |  |  |  |  |
|  | F-imd- | F-Δdpt | F-dpt<sup>S69R</sup> | F-dpt<sup>S69</sup> | M-imd- | M-Δdpt | M-dpt<sup>S69R</sup> |
| F-Δdpt | <u><2e-16</u> | - | - | - | - | - | - |
| F-dpt<sup>S69R</sup> | <u><2e-16</u> | <u>0.0348</u> | - | - | - | - | - |
| F-dpt<sup>S69</sup> | <u>4.00E-04</u> | <u>9.00E-04</u> | 0.9874 | - | - | - | - |
| M-imd- | <u><2e-16</u> | <u><2e-16</u> | <u><2e-16</u> | <u><2e-16</u> | - | - | - |
| M-Δdpt | <u>1.75E-02</u> | <u><2e-16</u> | 1.18E-01 | 6.87E-01 | <u><2e-16</u> | - | - |
| M-dpt<sup>S69R</sup> | <u>2.06E-02</u> | <u><2e-16</u> | 0.5139 | 9.63E-01 | <u><2e-16</u> | 0.9998 | - |
| M-dpt<sup>S69</sup> | 4.02E-01 | <u><2e-16</u> | <u>0.0333</u> | 2.97E-01 | <u><2e-16</u> | 0.9803 | 0.9241 |
| *A. tropicalis* |  |  |  |  |  |  |  |
|  | F-imd- | F-Δdpt | F-dpt<sup>S69R</sup> | F-dpt<sup>S69</sup> | M-imd- | M-Δdpt | M-dpt<sup>S69R</sup> |
| F-Δdpt | <u><2e-16</u> | - | - | - | - | - | - |
| F-dpt<sup>S69R</sup> | 0.2941 | <u><2e-16</u> | - | - | - | - | - |
| F-dpt<sup>S69</sup> | <u><2e-16</u> | 0.9003 | <u><2e-16</u> | - | - | - | - |
| M-imd- | <u>0.0016</u> | <u><2e-16</u> | 0.6689 | <u>0.0255</u> | - | - | - |
| M-Δdpt | <u>0.0017</u> | <u><2e-16</u> | 0.8677 | <u>4.00E-04</u> | 0.9987 | - | - |
| M-dpt<sup>S69R</sup> | 0.9522 | <u><2e-16</u> | 0.9472 | <u><2e-16</u> | 0.0876 | 0.1404 | - |
| M-dpt<sup>S69</sup> | 0.1403 | <u><2e-16</u> | 1 | <u>2.00E-04</u> | 0.8812 | 0.9821 | 0.8103 |
| *L. plantarum* + *P. rettgeri* |  |  |  |  |  |  |  |
|  | F-imd- | F-Δdpt | F-dpt<sup>S69R</sup> | F-dpt<sup>S69</sup> | M-imd- | M-Δdpt | M-dpt<sup>S69R</sup> |
| F-Δdpt | 0.1291 | - | - | - | - | - | - |
| F-dpt<sup>S69R</sup> | <u><2e-16</u> | <u><2e-16</u> | - | - | - | - | - |
| F-dpt<sup>S69</sup> | <u><2e-16</u> | <u><2e-16</u> | 0.3736 | - | - | - | - |
| M-imd- | <u><2e-16</u> | <u><2e-16</u> | <u>9.00E-04</u> | <u><2e-16</u> | - | - | - |
| M-Δdpt | <u><2e-16</u> | <u><2e-16</u> | <u>0.0176</u> | 0.9872 | <u><2e-16</u> | - | - |
| M-dpt<sup>S69R</sup> | <u><2e-16</u> | <u><2e-16</u> | 0.998 | 0.807 | <u>1.00E-04</u> | 0.1564 | - |
| M-dpt<sup>S69</sup> | <u><2e-16</u> | <u><2e-16</u> | 0.1446 | 0.9998 | <u><2e-16</u> | 1 | 0.5051 |
| *L. plantarum* + *A. tropicalis* |  |  |  |  |  |  |  |
|  | F-imd- | F-Δdpt | F-dpt<sup>S69R</sup> | F-dpt<sup>S69</sup> | M-imd- | M-Δdpt | M-dpt<sup>S69R</sup> |
| F-Δdpt | <u>0.008</u> | - | - | - | - | - | - |
| F-dpt<sup>S69R</sup> | <u><2e-16</u> | <u><2e-16</u> | - | - | - | - | - |
| F-dpt<sup>S69</sup> | <u><2e-16</u> | <u><2e-16</u> | 0.0632 | - | - | - | - |
| M-imd- | <u><2e-16</u> | <u><2e-16</u> | 0.9998 | 0.1957 | - | - | - |
| M-Δdpt | <u><2e-16</u> | <u><2e-16</u> | 0.9998 | 0.058 | 1 | - | - |
| M-dpt<sup>S69R</sup> | <u><2e-16</u> | <u><2e-16</u> | 0.8864 | 0.7167 | 0.9893 | 0.9573 | - |
| M-dpt<sup>S69</sup> | <u><2e-16</u> | <u><2e-16</u> | 0.6101 | 0.944 | 0.8792 | 0.7135 | 0.9997 |

**Table S5.**
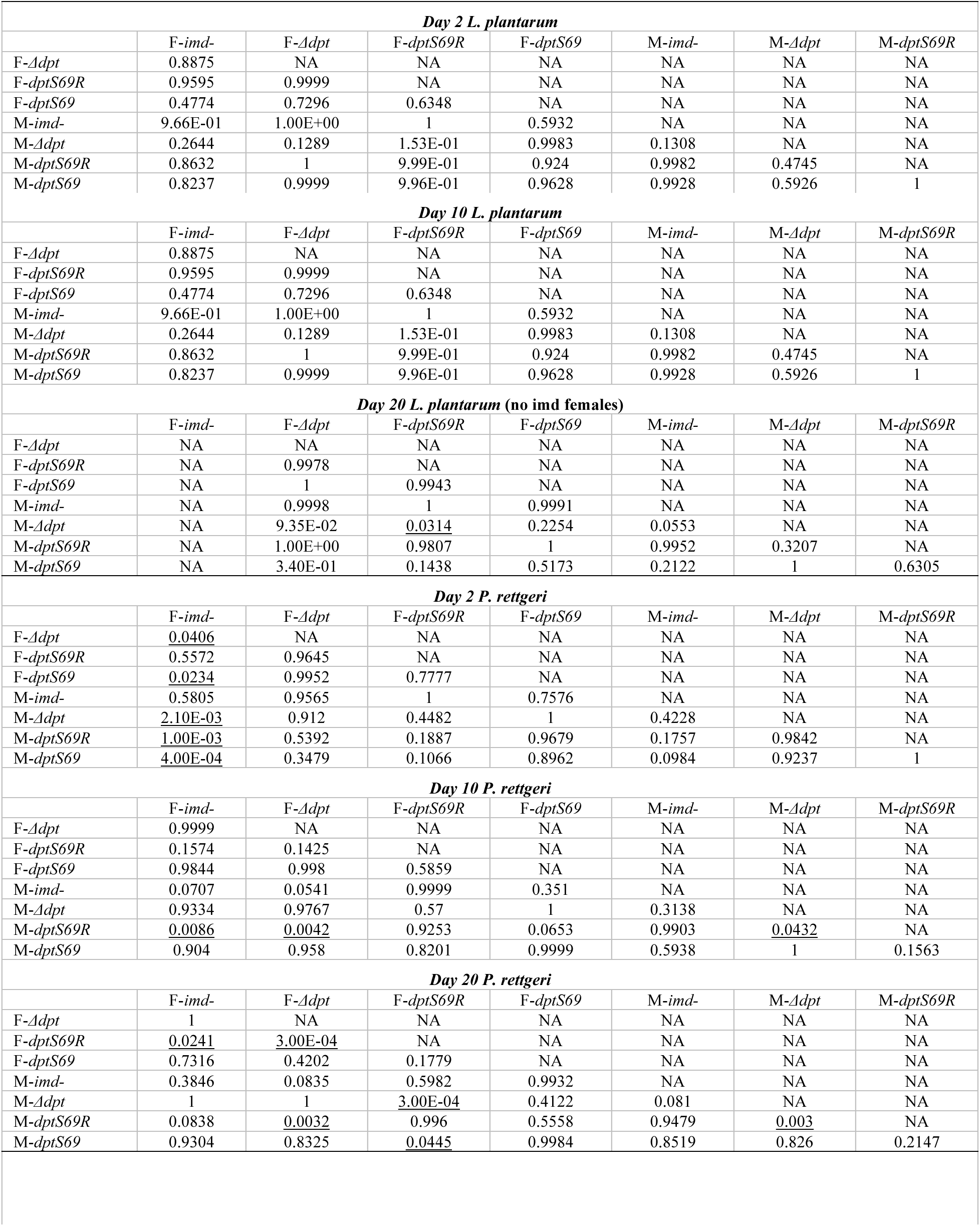
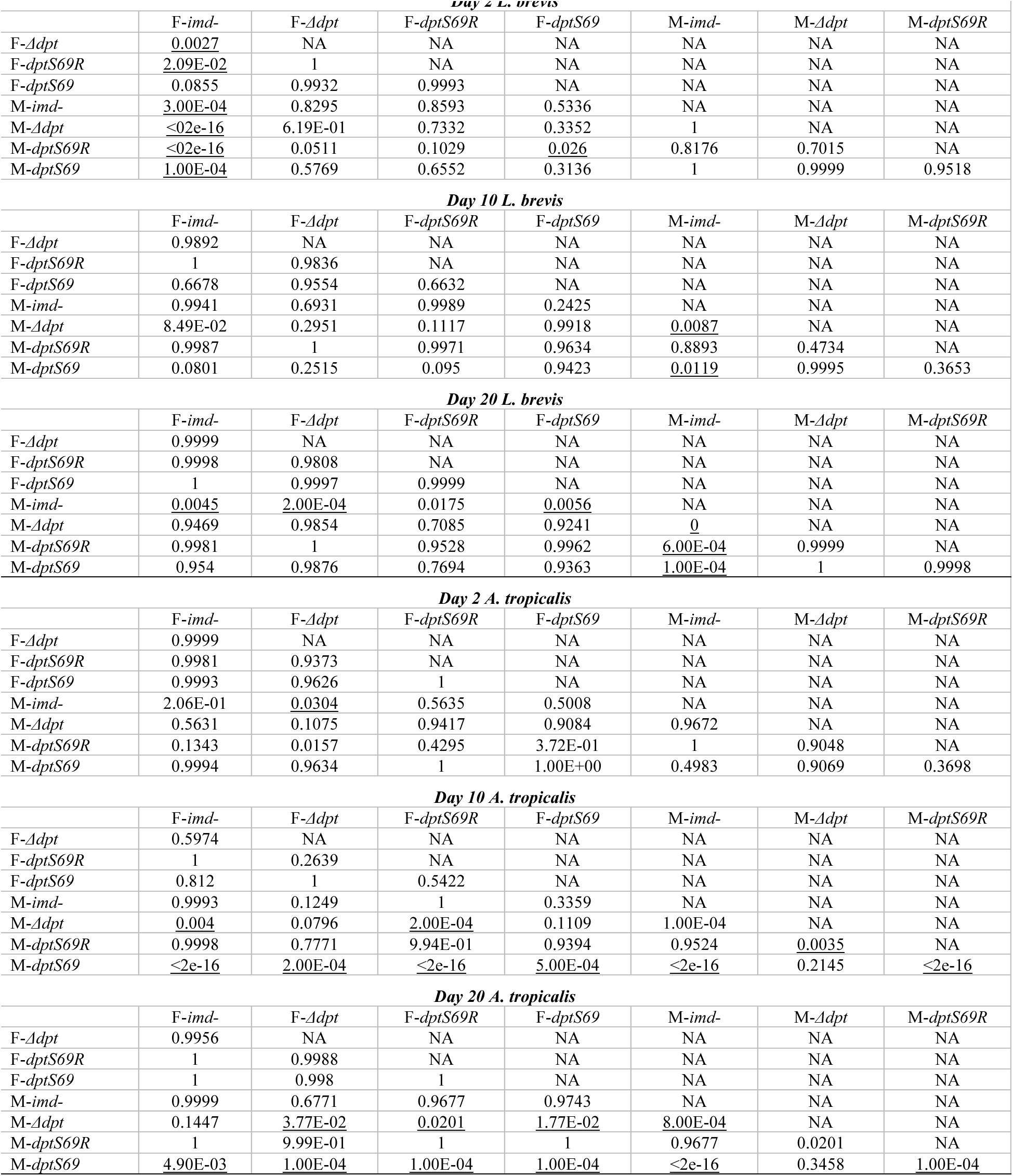
P-values from Tukey HSD for bacteria load. P-values less than 0.05 are underlined.

**Table S6.** P-values from Tukey HSD for bacterial load from poly-association with *L. plantarum* and *P. rettgeri*. P-values less than 0.05 are underlined.

| **Day 2 *L. plantarum*** |  |  |  |  |  |  |  |
| --- | --- | --- | --- | --- | --- | --- | --- |
|  | F-*imd*- | F-*Δdpt* | F-*dptS69R* | F-*dptS69* | M-*imd*- | M-*Δdpt* | M-*dptS69R* |
| F-*Δdpt* | 0.2226 | NA | NA | NA | NA | NA | NA |
| F-*dptS69R* | <u>0.0101</u> | 0.6004 | NA | NA | NA | NA | NA |
| F-*dptS69* | <u><2e-16</u> | <u>2.00E-04</u> | 0.1471 | NA | NA | NA | NA |
| M-*imd*- | 4.21E-01 | 1 | 0.7283 | <u>0.0017</u> | NA | NA | NA |
| M-*Δdpt* | <u>0.0031</u> | 4.83E-01 | 1 | <u>0.0344</u> | 0.6898 | NA | NA |
| M-*dptS69R* | <u><2e-16</u> | <u>6.00E-04</u> | 0.2442 | 1 | <u>0.0038</u> | 0.0725 | NA |
| M-*dptS69* | <u><2e-16</u> | <u>4.00E-04</u> | 0.2095 | 1 | <u>0.0029</u> | 0.0577 | 1 |
| **Day 10 *L. plantarum*** |  |  |  |  |  |  |  |
|  | F-*imd*- | F-*Δdpt* | F-*dptS69R* | F-*dptS69* | M-*imd*- | M-*Δdpt* | M-*dptS69R* |
| F-*Δdpt* | 0.2226 | NA | NA | NA | NA | NA | NA |
| F-*dptS69R* | <u>0.0101</u> | 0.6004 | NA | NA | NA | NA | NA |
| F-*dptS69* | <u><2e-16</u> | <u>2.00E-04</u> | 0.1471 | NA | NA | NA | NA |
| M-*imd*- | 4.21E-01 | 1 | 0.7283 | <u>0.0017</u> | NA | NA | NA |
| M-*Δdpt* | 0.0031 | 4.83E-01 | 1 | <u>0.0344</u> | 0.6898 | NA | NA |
| M-*dptS69R* | <u><2e-16</u> | <u>6.00E-04</u> | 0.2442 | 1 | <u>0.0038</u> | 0.0725 | NA |
| M-*dptS69* | <u><2e-16</u> | <u>4.00E-04</u> | 0.2095 | 1 | <u>0.0029</u> | 0.0577 | 1 |
| **Day 20 *L. plantarum* (no imd females)** |  |  |  |  |  |  |  |
|  | F-*imd*- | F-*Δdpt* | F-*dptS69R* | F-*dptS69* | M-*imd*- | M-*Δdpt* | M-*dptS69R* |
| F-*Δdpt* | NA | NA | NA | NA | NA | NA | NA |
| F-*dptS69R* | NA | 0.831 | NA | NA | NA | NA | NA |
| F-*dptS69* | NA | 0.9883 | 0.9977 | NA | NA | NA | NA |
| M-*imd*- | NA | 0.5334 | 0.9998 | 0.9429 | NA | NA | NA |
| M-*Δdpt* | NA | 0.9999 | 0.484 | 0.857 | 0.1772 | NA | NA |
| M-*dptS69R* | NA | 0.9202 | 1 | 0.9999 | 0.9943 | 0.6188 | NA |
| M-*dptS69* | NA | 0.9999 | 0.5746 | 0.8987 | 0.2615 | 1 | 0.7064 |
| **Day 2 *P. rettgeri*** |  |  |  |  |  |  |  |
|  | F-*imd*- | F-*Δdpt* | F-*dptS69R* | F-*dptS69* | M-*imd*- | M-*Δdpt* | M-*dptS69R* |
| F-*Δdpt* | 0.0784 | NA | NA | NA | NA | NA | NA |
| F-*dptS69R* | 0.1199 | 1 | NA | NA | NA | NA | NA |
| F-*dptS69* | 0.434 | 0.9992 | 0.9966 | NA | NA | NA | NA |
| M-*imd*- | 0.8069 | 0.912 | 0.8957 | 0.9988 | NA | NA | NA |
| M-*Δdpt* | 0.0063 | 0.9457 | 0.9972 | 0.8097 | 0.3815 | NA | NA |
| M-*dptS69R* | 0.131 | 1 | 1 | 0.9977 | 0.9101 | 0.9957 | NA |
| M-*dptS69* | <u>0.0348</u> | 0.9902 | 0.9996 | 0.9254 | 0.6186 | 1 | 0.9993 |
| **Day 10 *P. rettgeri*** |  |  |  |  |  |  |  |
|  | F-*imd*- | F-*Δdpt* | F-*dptS69R* | F-*dptS69* | M-*imd*- | M-*Δdpt* | M-*dptS69R* |
| F-*Δdpt* | 1 | NA | NA | NA | NA | NA | NA |
| F-*dptS69R* | 0.993 | 0.9967 | NA | NA | NA | NA | NA |
| F-*dptS69* | 0.9835 | 0.9901 | 1 | NA | NA | NA | NA |
| M-*imd*- | 0.998 | 0.9994 | 1 | 1 | NA | NA | NA |
| M-*Δdpt* | 1 | 0.9997 | 0.9485 | 0.9085 | 0.9773 | NA | NA |
| M-*dptS69R* | 1 | 1 | 0.9998 | 0.9992 | 1 | 0.9981 | NA |
| M-*dptS69* | 0.9911 | 0.9954 | 1 | 1 | 1 | 0.9393 | 0.9998 |
| **Day 20 *P. rettgeri* (no imd females)** |  |  |  |  |  |  |  |
|  | F-*imd*- | F-*Δdpt* | F-*dptS69R* | F-*dptS69* | M-*imd*- | M-*Δdpt* | M-*dptS69R* |
| F-*Δdpt* | NA | NA | NA | NA | NA | NA | NA |
| F-*dptS69R* | NA | 0.9927 | NA | NA | NA | NA | NA |
| F-*dptS69* | NA | 0.9971 | 0.8027 | NA | NA | NA | NA |
| M-*imd*- | NA | 0.9999 | 0.911 | 1 | NA | NA | NA |
| M-*Δdpt* | NA | 9.64E-01 | 1 | 0.5715 | 0.7503 | NA | NA |
| M-*dptS69R* | NA | 1 | 0.9896 | 0.9966 | 0.9998 | 0.9478 | NA |
| M-*dptS69* | NA | 0.9991 | 1 | 0.8929 | 0.9652 | 0.9998 | 0.9986 |

**Table S7.** Desiccation resistance Tukey HSD p-values. P-values less than 0.05 are underlined. “A” refers to flies reared axenically, “C” refers to flies reared on a conventional diet.

|  | A<br>*dptS69* | A<br>*Δdpt* | A<br>*dptS69R* | A<br>*imd-* | A<br>OreR | A<br>w1118 | C<br>*dptS69* | C<br>*Δdpt* | C<br>*dptS69R* | C<br>*imd-* | C<br>OreR |
| --- | --- | --- | --- | --- | --- | --- | --- | --- | --- | --- | --- |
| A *Δdpt* | 0.9999 | NA | NA | NA | NA | NA | NA | NA | NA | NA | NA |
| A *dptS69R* | 0.3541 | 0.9978 | NA | NA | NA | NA | NA | NA | NA | NA | NA |
| A *imd-* | 0.9195 | 0.8332 | <u>0.003</u> | NA | NA | NA | NA | NA | NA | NA | NA |
| A OreR | 1 | 1 | 0.5898 | 0.8561 | NA | NA | NA | NA | NA | NA | NA |
| A w1118 | <u><2e-16</u> | <u>7.00E-04</u> | <u><2e-16</u> | <u>0.007</u> | <u><2e-16</u> | NA | NA | NA | NA | NA | NA |
| C *dptS69* | <u><2e-16</u> | 0.1208 | 0.1473 | <u><2e-16</u> | <u>1.00E-04</u> | <u><2e-16</u> | NA | NA | NA | NA | NA |
| C *Δdpt* | <u><2e-16</u> | <u>5.00E-04</u> | <u>2.00E-04</u> | <u><2e-16</u> | <u><2e-16</u> | <u><2e-16</u> | 0.3672 | NA | NA | NA | NA |
| C *dptS69R* | <u><2e-16</u> | <u>0.0193</u> | <u>0.0086</u> | <u><2e-16</u> | <u><2e-16</u> | <u><2e-16</u> | 0.9995 | 0.8062 | NA | NA | NA |
| C *imd-* | <u>0.0057</u> | 0.7281 | 0.963 | <u><2e-16</u> | <u>0.0242</u> | <u><2e-16</u> | 0.934 | <u>0.013</u> | 0.3953 | NA | NA |
| C OreR | <u>0.0024</u> | 0.5452 | 0.8351 | <u><2e-16</u> | <u>0.0105</u> | <u><2e-16</u> | 0.9979 | 0.0653 | 0.8078 | 1 | NA |
| C w1118 | 0.9996 | 1 | 0.8729 | 0.3974 | 1 | <u><2e-16</u> | <u>3.00E-04</u> | <u><2e-16</u> | <u><2e-16</u> | 0.0781 | <u>0.0352</u> |

**Table S8.**
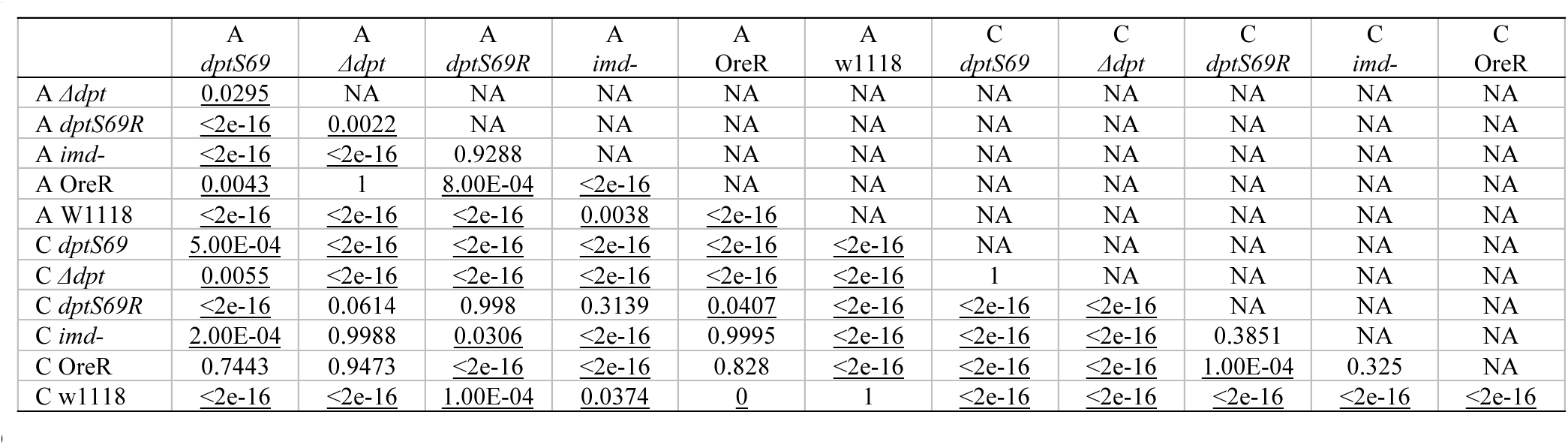
Starvation resistance Tukey HSD p-values. P-values less than 0.05 are underlined. “A” refers to flies reared axenically, “C” refers to flies reared on a conventional diet.

**Table S9.** Conventional longevity Cox proportional hazards regression p-values. P-values less than 0.05 are underlined. “F” and “M” refer to males and females, respectively; “het” refers to heterozygotes for serine/arginine.

|  | F<br>het | F<br>*imd-* | F<br>*Δdpt* | F<br>*dptS69R* | F<br>*dptS69* | M<br>het | M<br>*imd-* | M<br>*Δdpt* | M<br>*dptS69R* |
| --- | --- | --- | --- | --- | --- | --- | --- | --- | --- |
| F *imd-* | 1 | NA | NA | NA | NA | NA | NA | NA | NA |
| F *Δdpt* | 1 | 1 | NA | NA | NA | NA | NA | NA | NA |
| F *dptS69R* | <u><2e-16</u> | <u><2e-16</u> | <u><2e-16</u> | NA | NA | NA | NA | NA | NA |
| F *dptS69* | <u><2e-16</u> | <u>3.00E-04</u> | <u><2e-16</u> | 0.6785 | NA | NA | NA | NA | NA |
| M het | <u><2e-16</u> | <u><2e-16</u> | <u><2e-16</u> | 0.9797 | <u>0.0443</u> | NA | NA | NA | NA |
| M *imd-* | 0.1346 | 0.3182 | <u>0.0498</u> | <u>0.0025</u> | 0.5097 | 0 | NA | NA | NA |
| M *Δdpt* | <u><2e-16</u> | <u><2e-16</u> | <u><2e-16</u> | 0.948 | 0.9977 | 0.1163 | <u>0.0321</u> | NA | NA |
| M *dptS69R* | <u><2e-16</u> | <u><2e-16</u> | <u><2e-16</u> | 0.0622 | <u><2e-16</u> | 0.4311 | <u><2e-16</u> | <u><2e-16</u> | NA |
| M *dptS69* | <u><2e-16</u> | <u><2e-16</u> | <u><2e-16</u> | 0.9984 | 0.9882 | 0.5841 | <u>0.0492</u> | 1 | <u>0.0043</u> |

**Table S10.**
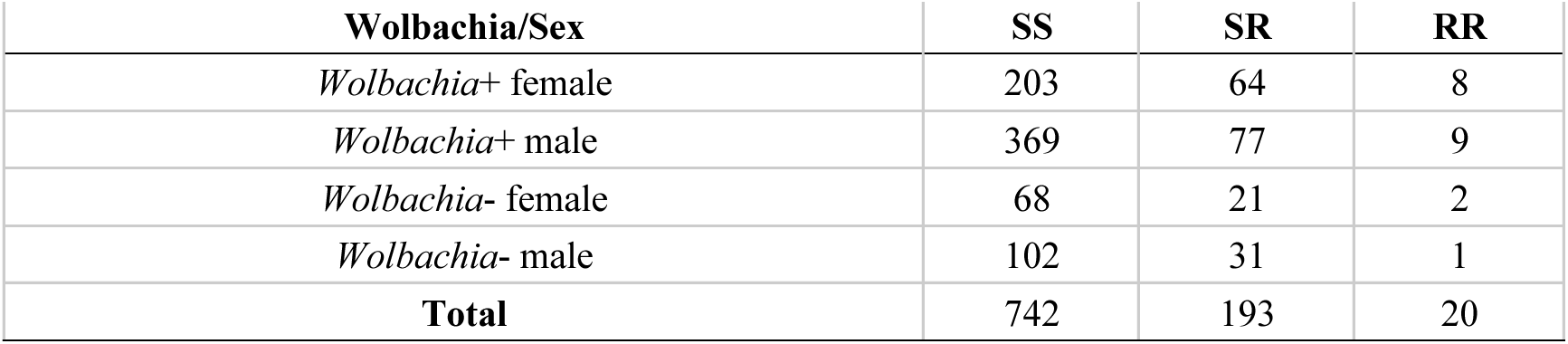
Genotyped wild caught *D. melanogaster*.

## Supplemental Figures

**Figure S1.**
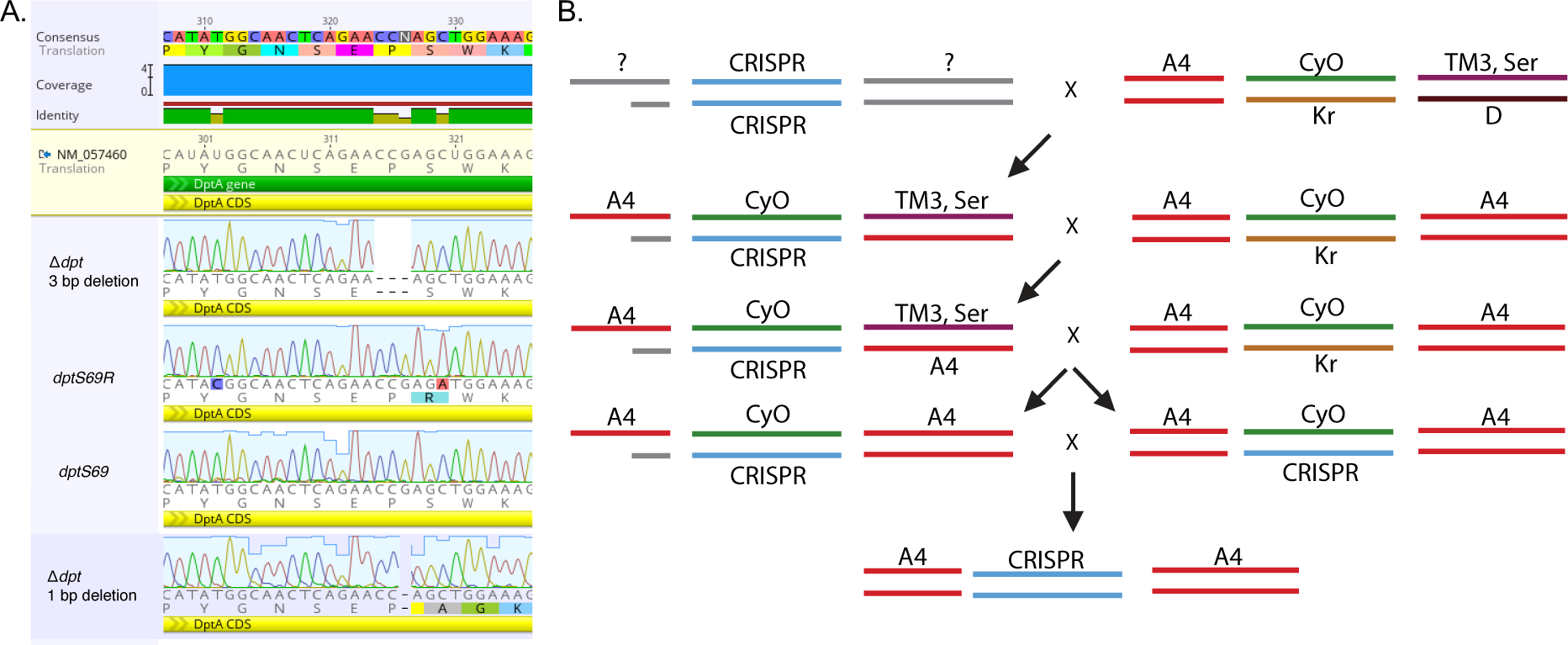
CRISPR/Cas9 genome editing of the *dpt* locus. A) Alignment of CRISPR/Cas9 genome edited lines. CRISPR/Cas9 genome edited lines aligned to consensus sequence. Genotype/line name is indicated on the left-hand side of the figure. B) Crossing schematic for CRISPR clean up. A modified Bloomington stock # 7198 was used for CRISPR injections. The line contained *CyO*/*Kruppel* balanced on the 2^nd^ chromosome, and *serrate*/*Dichaete* balanced on the 3^rd^ chromosome. Our version, 7198^A4^, was provided by Stuart Macdonald and has the DSPR (King et al. 2012) A4 line’s X chromosome instead of the *w*[*] from the original 7198.

**Figure S2.**
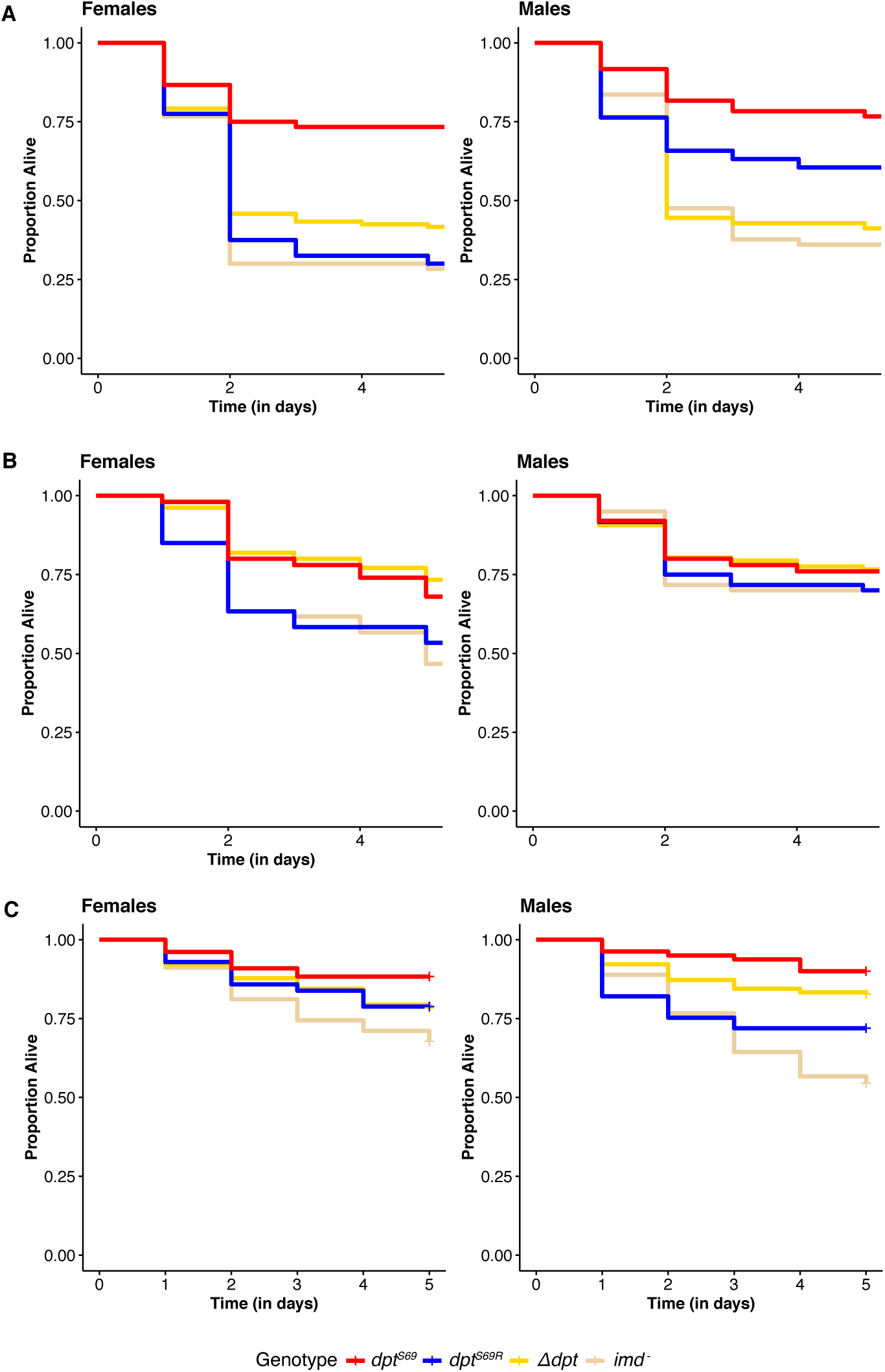
Male vs. female survival after systemic infection. Systemic infections were repeated with females and males. A) *P. rettgeri* and B) *E. faecalis* are normal systemic infection bacteria in *Drosophila*. C) *L. plantarum* was used to test differences in systemic based on sex based on the results from gnotobiotic longevity.

**Figure S3.**
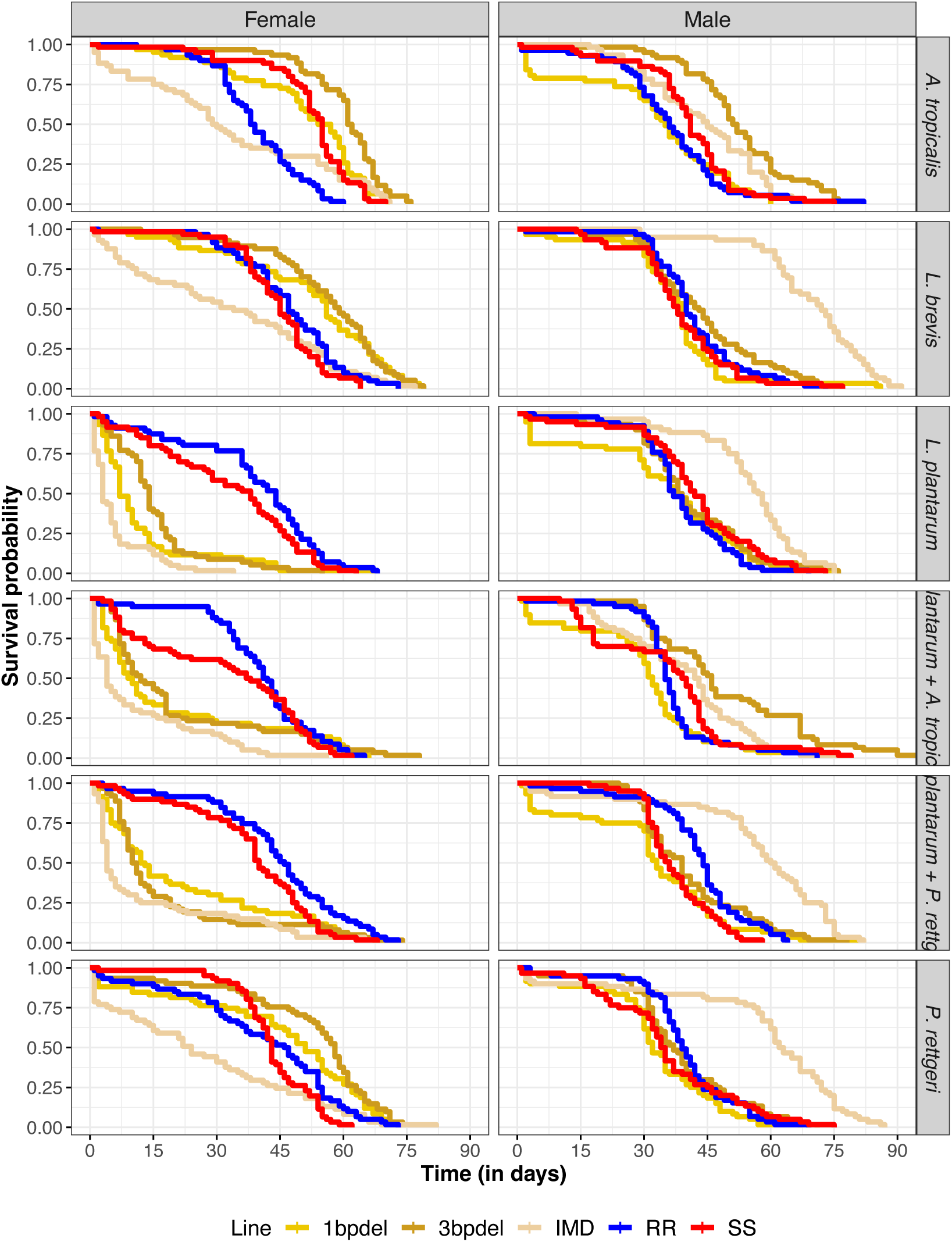
Gnotobiotic longevity. Survival plots of all bacteria. Dpt null lines have been separated into the two lines used, 1 base pair deletion and 3 bp deletion.

**Figure S4.**
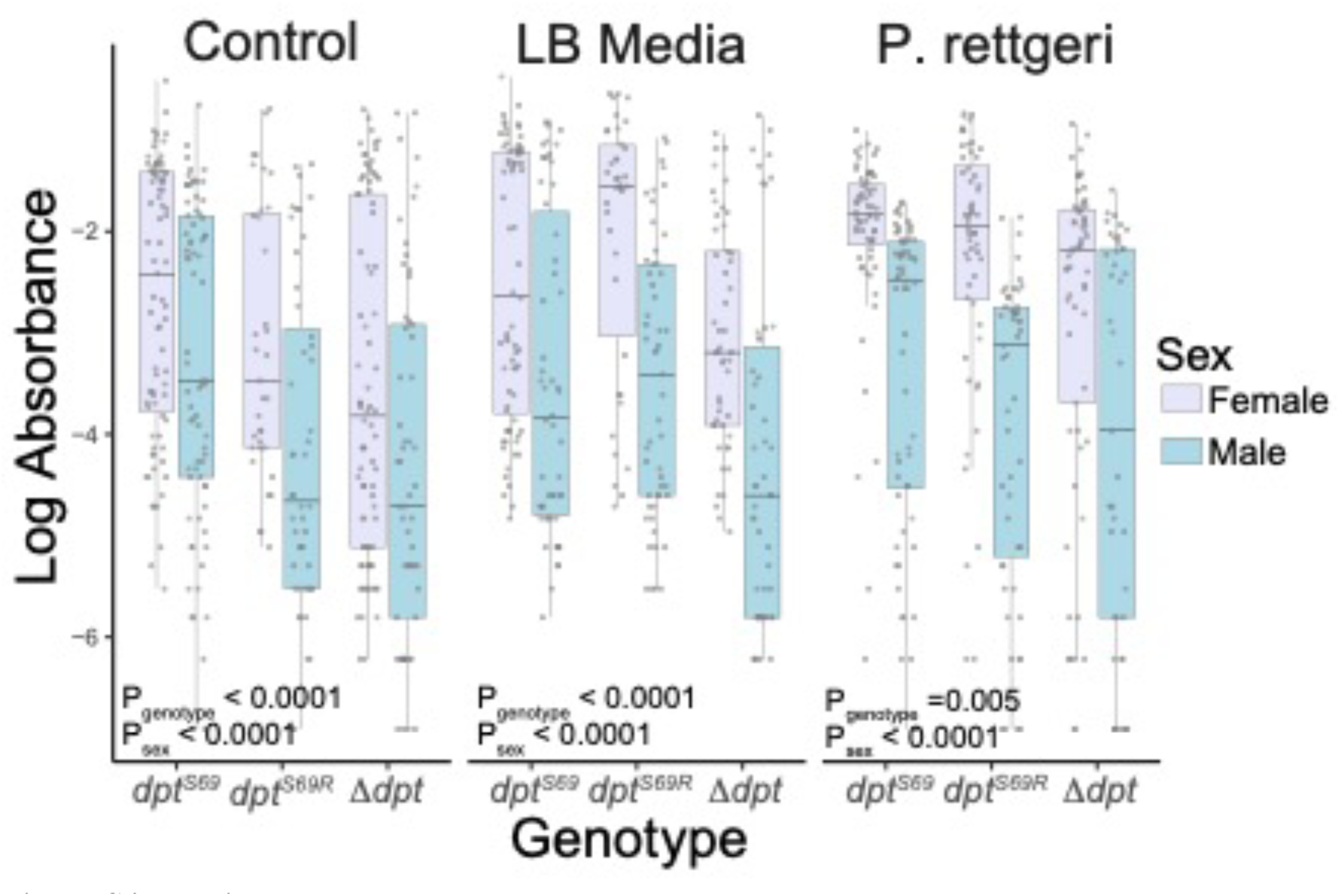
Feeding Rates. Female flies eat more food (as measured by blue dye ingestion) than males, but genotype also influences feeding. Each genotype has more than 100 individuals per treatment per sex. Control refers to no added media, LB Media is the addition of sterile MB, and *P. rettgeri* is the addition of *P. rettgeri* at a concentration of OD600 = 15 (±1). Absorbance was measured at 630nm.

**Figure S5.**
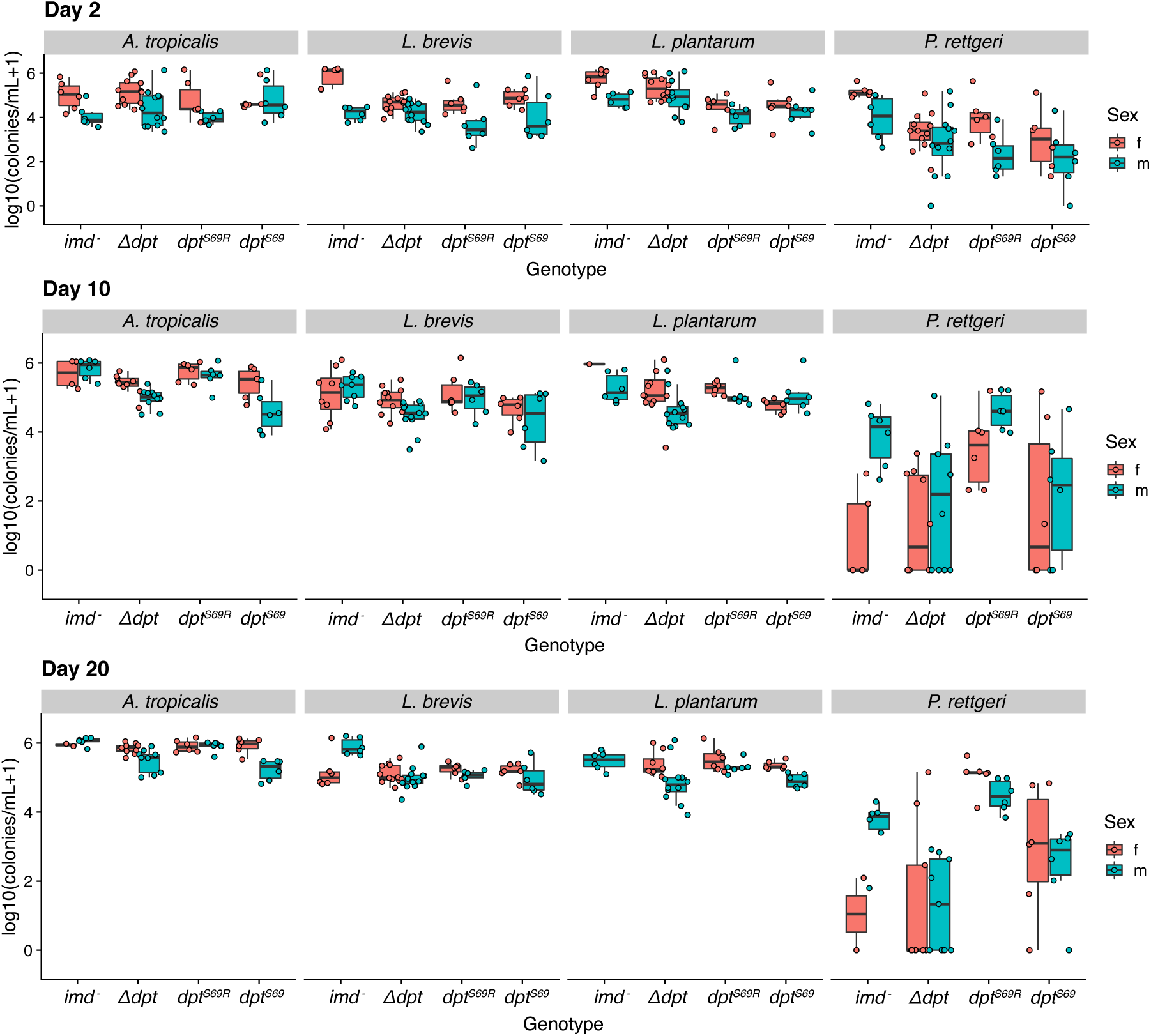
Bacterial load at day 2-, 10-, and 20-days post feeding. All bacteria were plated on MRS media except for *P. rettgeri* that was plated on LB media to grow colonies. Significance can be found in Table 2.S6. Individual points represent a pool of 3 flies. There is no *imd-* female bar for *L. plantarum* on day 20 because all female flies had died at this timepoint.

**Figure S6.**
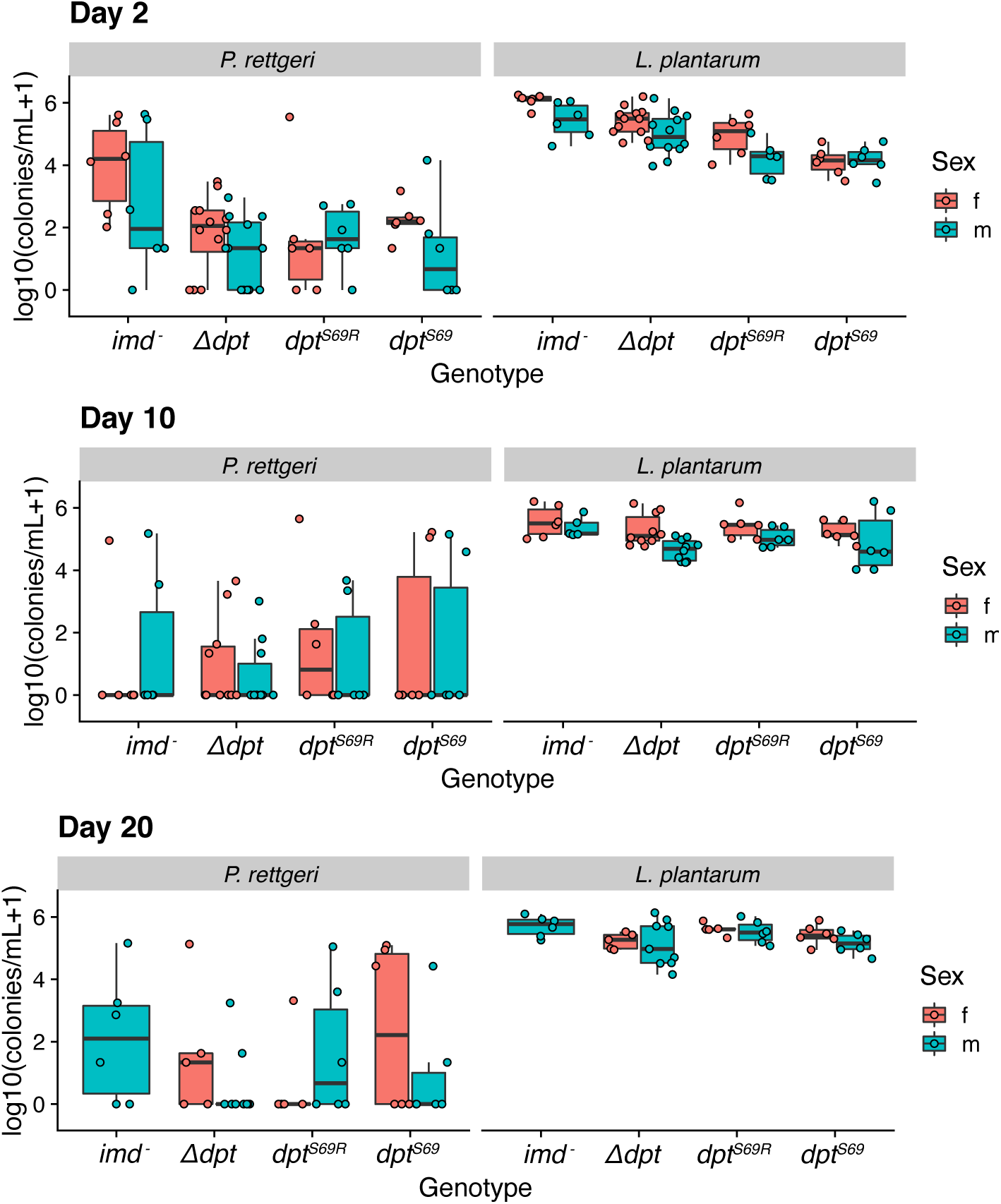
Bacterial load of *L. plantarum* and *P. rettgeri* poly-association. The flies were co-feed with *L. plantarum* and *P. rettgeri* and plated on MRS and LB respectively to select for the different bacteria. There is no *imd-* female bar on day 20 because all flies had died at this timepoint.

**Figure S7.**
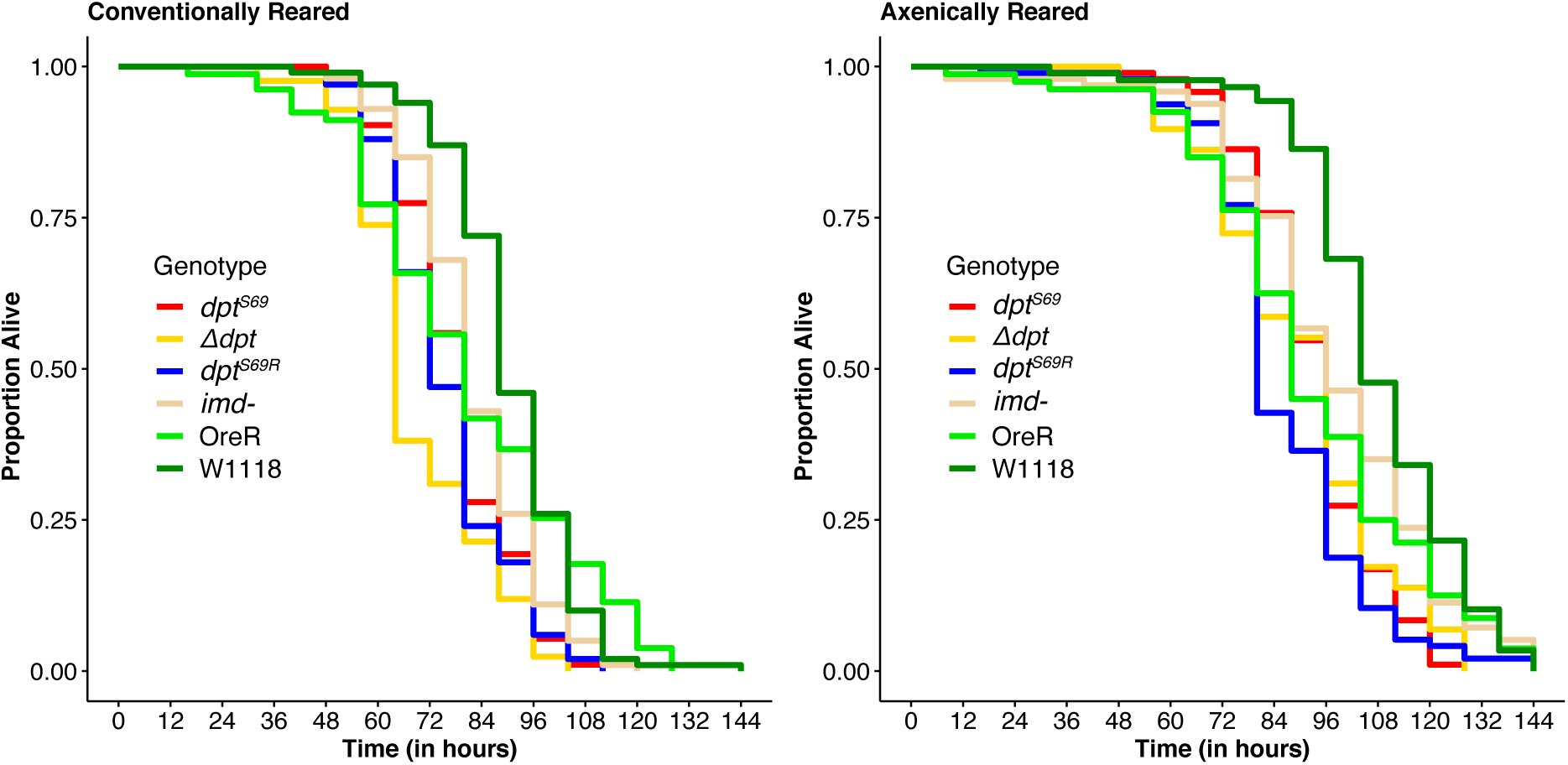
Desiccation stress survival with additional wildtype lines (OreR and W1118) OreR and W1118 are standard *Drosophila melanogaster* lab stocks. They are both homozygous serine but have different resistance to desiccation stress than *dpt^S69^*.

**Figure S8.**
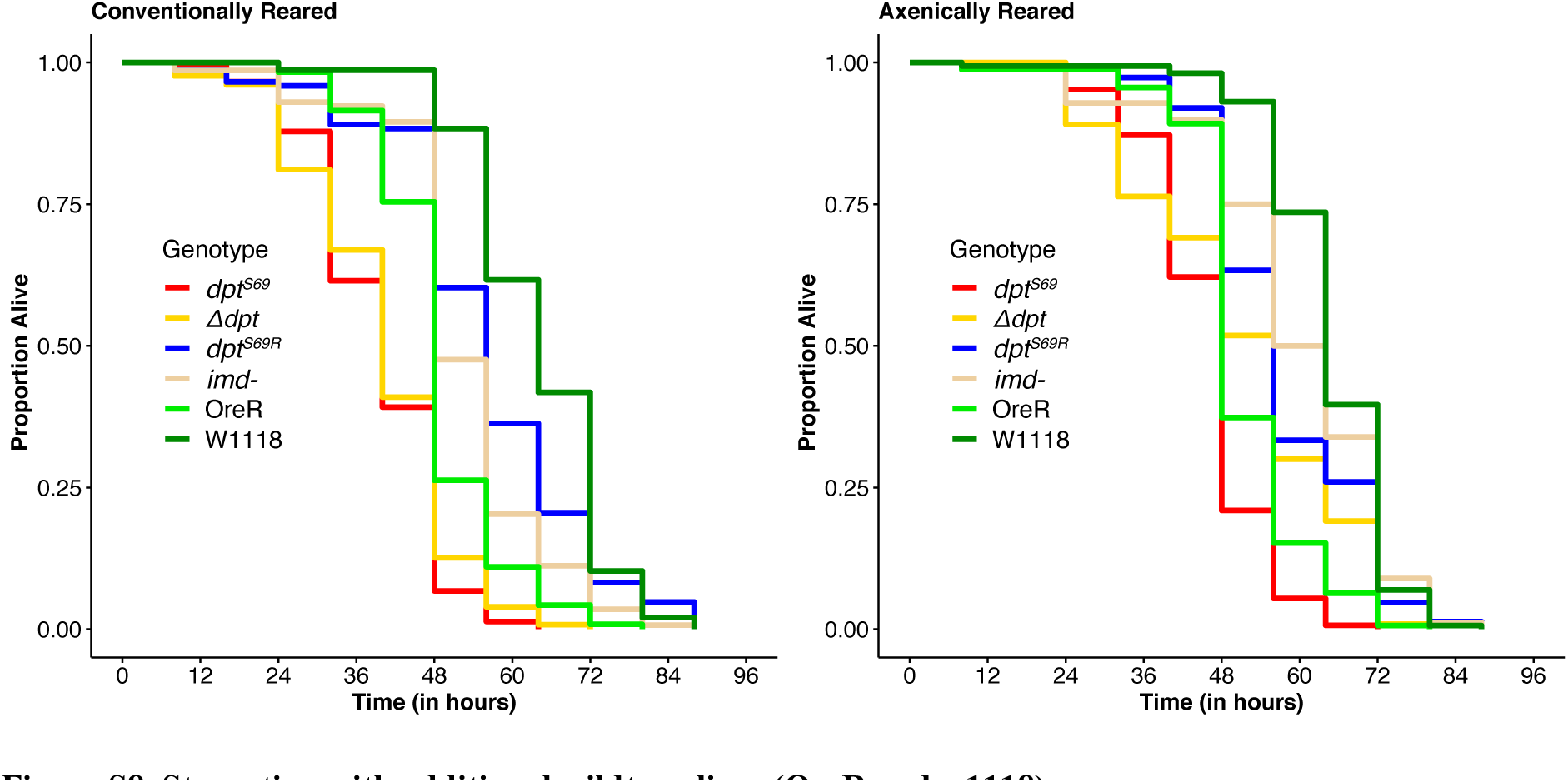
Starvation with additional wildtype lines (OreR and w1118) OreR and W1118 are standard *Drosophila melanogaster* lab stocks. They are both homozygous serine but have different resistance to starvation stress than *dpt^S69^*.

**Figure S9.**
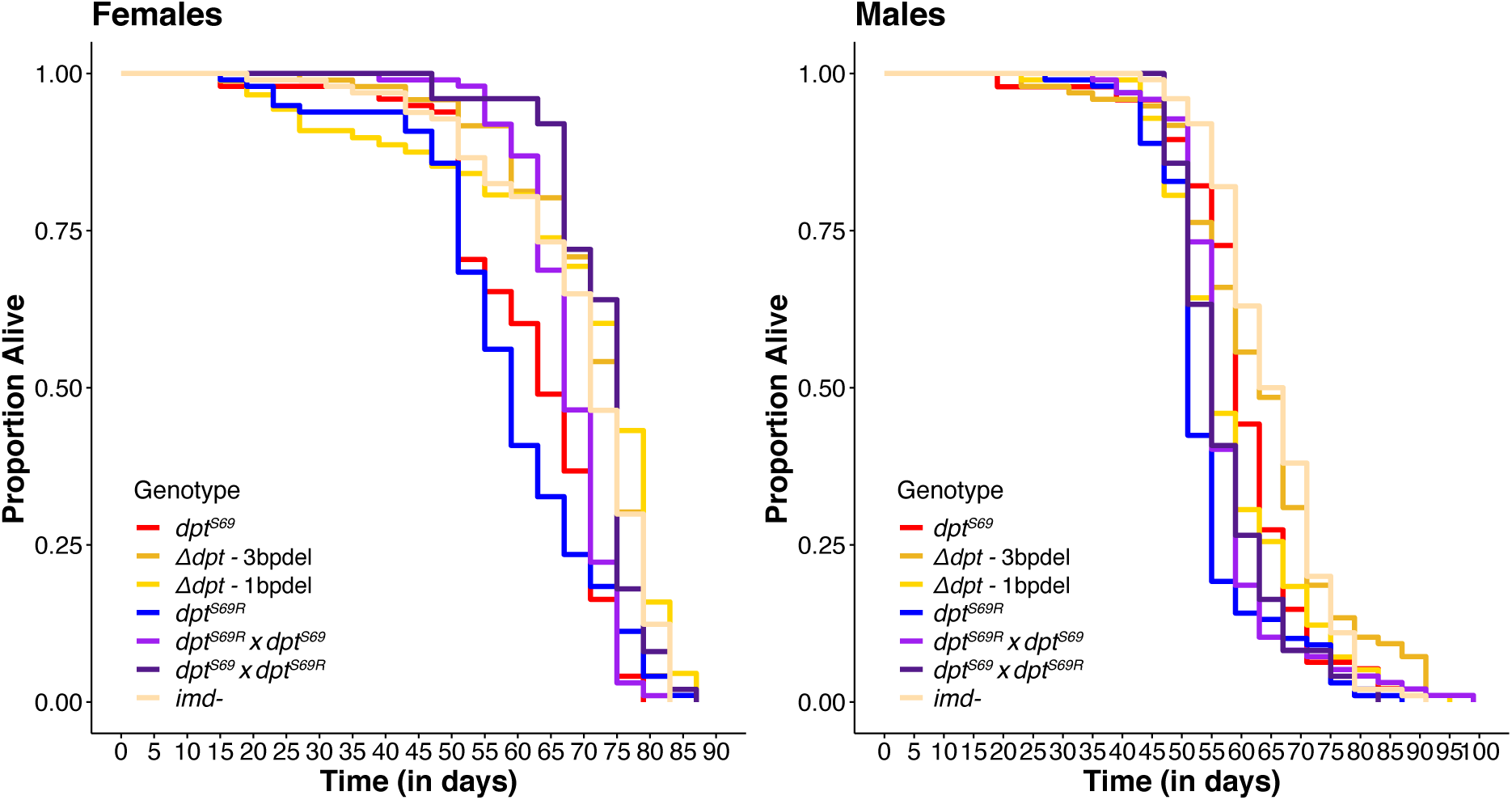
Longevity with separated *Δdpt* lines and heterozygotes. Two *Δdpt* lines were used. One with a 1 base pair deletion and the other with a 3 base pair deletion. These lines were combined for ease of viewing in Fig 4C. Heterozygotes were made with reciprocal crosses

## Notes

### Competing Interest Statement

The authors have declared no competing interest.

### Summary of Updates

This revision has been revised to address reviewer comments as part of eLife's reviewed preprints program.

